# Synapse type-specific proteomic dissection identifies IgSF8 as a hippocampal CA3 microcircuit organizer

**DOI:** 10.1101/846816

**Authors:** Nuno Apóstolo, Samuel N. Smukowski, Jeroen Vanderlinden, Giuseppe Condomitti, Vasily Rybakin, Jolijn ten Bos, Laura Trobiani, Sybren Portegies, Kristel M. Vennekens, Natalia V. Gounko, Davide Comoletti, Keimpe D. Wierda, Jeffrey N. Savas, Joris de Wit

## Abstract

Synaptic diversity is a key feature of neural circuits. The structural and functional diversity of closely spaced inputs converging on the same neuron suggests that cell-surface interactions are essential in organizing input properties. Here, we analyzed the cell-surface protein (CSP) composition of hippocampal mossy fiber (MF) inputs on CA3 pyramidal neurons to identify regulators of MF-CA3 synapse properties. We uncover a rich cell-surface repertoire that includes adhesion proteins, guidance cue receptors, extracellular matrix (ECM) proteins, and uncharacterized CSPs. Interactome screening reveals multiple ligand-receptor modules and identifies ECM protein Tenascin-R (TenR) as a ligand of the uncharacterized neuronal receptor IgSF8. Presynaptic *Igsf8* deletion impairs MF-CA3 synaptic architecture and robustly decreases the density of bouton filopodia that provide feedforward inhibition of CA3 neurons. Consequently, loss of IgSF8 increases CA3 neuron excitability. Our findings identify IgSF8 as a regulator of CA3 microcircuit development and suggest that combinations of CSP modules define input identity.

## Introduction

Neural circuits are composed of distinct neuronal cell types connected in highly specific patterns. Large pyramidal neurons integrate information from multiple classes of presynaptic neurons that form different types of synapses on their dendrites. These synapses have a distinct subcellular location, architecture, and functional properties (O’Rourke et al., 2012), all of which affect information processing in pyramidal neurons (Spruston, 2008).

The hippocampal CA3 circuit offers a striking example of input diversification. CA3 pyramidal neurons receive excitatory input on their proximal apical dendrite from dentate gyrus (DG) granule cell (GC) axons, or mossy fibers (MFs), in stratum lucidum (SL) (Figure 1A). MF boutons are unusually large, contain multiple release sites, and have filopodia that extend from the main bouton to form excitatory synapses on nearby interneurons (Acsády et al., 1998). These filopodia provide feedforward inhibition of CA3 neurons, important for the control of CA3 excitability (Martin et al., 2015; Torborg et al., 2010) and memory precision (Guo et al., 2018; Ruediger et al., 2011). The postsynaptic compartment of the MF-CA3 synapse consists of multi-headed dendritic spines, or thorny excrescences, that are engulfed by the presynaptic bouton (Chicurel and Harris, 1992; Rollenhagen and Lübke, 2010; Wilke et al., 2013). Compared to the MF inputs, recurrent CA3 inputs on the medial portion of the CA3 apical dendrite in stratum radiatum are much smaller and morphologically less elaborate (Nicoll and Schmitz, 2005). Short-term synaptic plasticity differs markedly at these two types of excitatory inputs (Nicoll and Schmitz, 2005; Salin et al., 1996), which is important for memory encoding (Rebola et al., 2017). The molecular mechanisms organizing these input-specific properties remain poorly understood.

**Figure 1.**
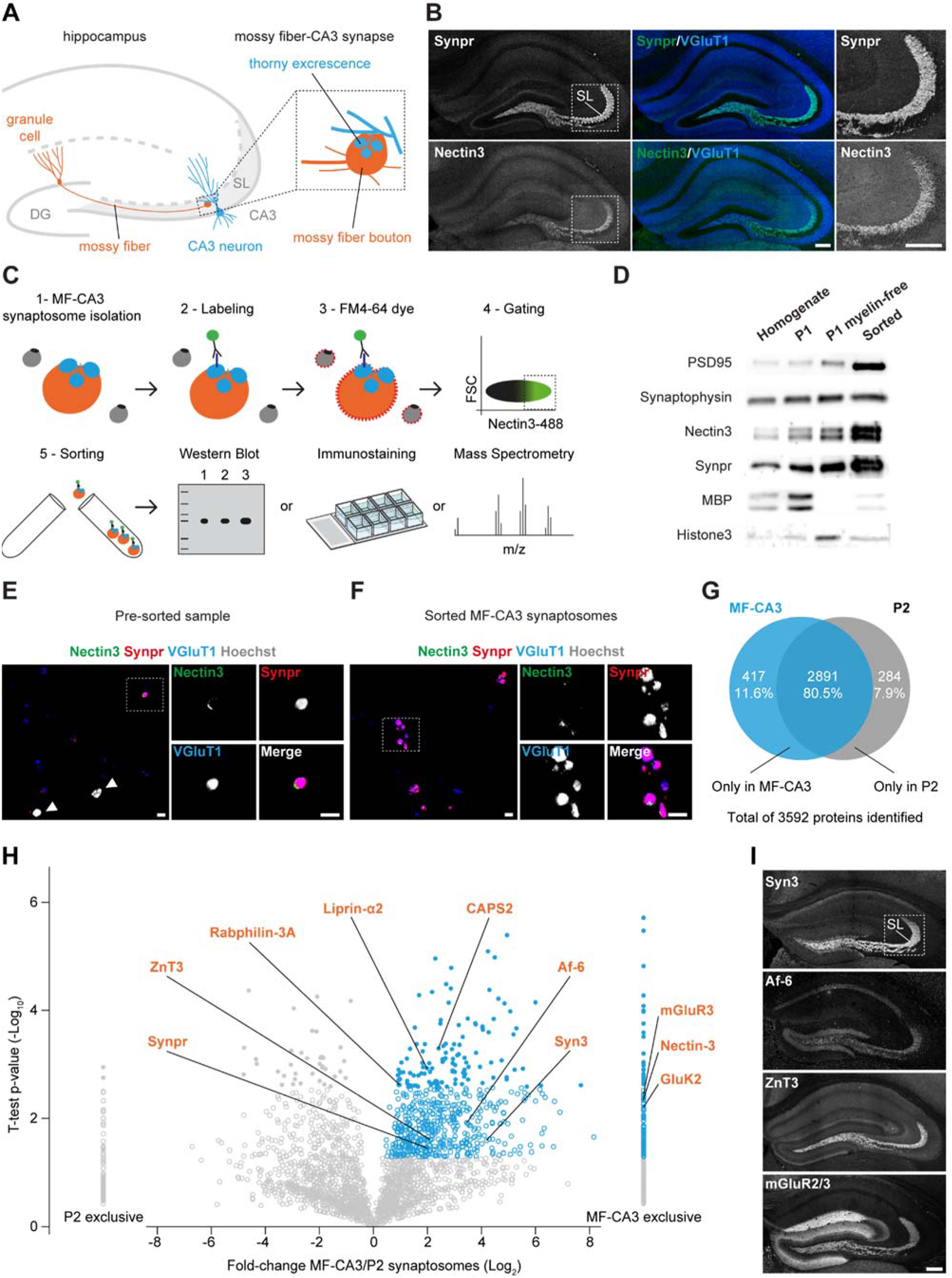
Isolation of MF-CA3 synaptosomes. (**A**) Cartoon illustrating the large hippocampal MF-CA3 synapse. (**B**) Confocal images of P28 mouse hippocampal sections immunostained for Synpr, Nectin-3 and VGluT1. Magnified insets of SL in CA3 are shown on the right. (**C**) Workflow to isolate and analyze MF-CA3 synaptosomes. (**D**) Validation of enrichment for MF-CA3 synaptosomes in sorted material by western blot. MPB, Myelin-binding protein. (**E**) and (**F**) Confocal images of pre-sorted and sorted material, respectively, immunostained for Nectin-3, Synpr, VGluT1 and Hoechst. (**G**) Venn diagram capturing number and distribution of proteins identified in sorted MF-CA3 synaptosomes and P2 synaptosomes by LC-MS/MS in three independent experiments (10-12 mice per experiment). (**H**) Relative distribution of proteins detected in sorted MF-CA3 synaptosomes and P2 synaptosomes. Significant proteins with positive MF-CA3/P2 log_2_ fold-change are highlighted in blue (p-value ≤ 0.05, Student’s t-test). High-confidence measurements at a 5% FDR are shown as closed circles (q-value ≤ 0.05, Benjamini-Hochberg correction). A selection of known MF-CA3 synaptic proteins is annotated in orange. (**I**) Confocal images of P28 mouse hippocampal sections immunostained for known MF-CA3 synaptic markers detected in sorted MF-CA3 synaptosomes. Scale bars in (B) and (I) 200 μm, in (E) and (F) 5 μm. See also Figure S1.

The striking diversity of neighboring excitatory inputs on the same CA3 dendrite suggests that short-range, cell-surface interactions play an important role in regulating input-specific properties. Cell-surface proteins (CSPs), including transmembrane, membrane-anchored, and secreted proteins, are expressed in cell type-specific combinations (Földy et al., 2016; Li et al., 2017; Paul et al., 2017; Shekhar et al., 2016) and form protein-protein interaction networks to control neurite guidance, target selection, and synapse development required for the formation of functional circuits (Sanes and Yamagata, 2009; Shen and Scheiffele, 2010; de Wit and Ghosh, 2016). Recent studies reveal an input-specific localization and function of several postsynaptic CSPs in hippocampal CA3 and CA1 pyramidal neuron dendrites (Condomitti et al., 2018; DeNardo et al., 2012; Sando et al., 2019; Schroeder et al., 2018). Distinct classes of interneurons innervating different domains of cortical pyramidal neurons express cell type-specific CSPs that regulate domain-specific synapse formation (Favuzzi et al., 2019). These studies suggest that compartmentalized distributions of CSPs in dendrites and differential expression in axonal populations contribute to defining input-specific properties (Apóstolo and de Wit, 2019), but analyzing CSP composition of specific inputs remains challenging.

Here, we isolate MF-CA3 synaptosomes and use mass spectrometry (MS) to begin uncovering their CSP composition and identify novel regulators of MF-CA3 synapse properties. We identify a rich CSP repertoire that includes adhesion proteins, guidance cue receptors, secreted and ECM proteins, as well as several uncharacterized CSPs. Approximately 80% of these CSPs has not been reported to localize or function at MF-CA3 synapses and approximately 50% does not have an annotated synaptic function. Interactome screening reveals multiple ligand-receptor modules among these CSPs and identifies the secreted ECM protein Tenascin-R (TenR) as a novel ligand of IgSF8, an uncharacterized neuronal receptor strongly enriched in the MF-CA3 pathway. Dentate GC-specific deletion of *Igsf8* reduces the number of release sites in MF boutons and robustly decreases the number of filopodia. Functional analysis reveals reduced feedforward inhibition and increased excitability of CA3 pyramidal neurons in the absence of presynaptic IgSF8. Together, our findings reveal a diverse CSP composition of MF-CA3 synapses, identify IgSF8 as a regulator of CA3 microcircuit connectivity and function, and suggest that combinations of CSP modules specify MF-CA3 synapse identity.

## Results

### Isolation of MF-CA3 synaptosomes

To isolate MF-CA3 synapses and start uncovering their CSP composition, we relied on two key features of the MF-CA3 synapse: its large size and the presence of puncta adherentia (PA), a specialized type of adhesive junction found at large synapses (Rollenhagen and Lübke, 2006) that is morphologically and molecularly distinct from the synaptic junction. We combined two established approaches to take advantage of these features. First, we prepared MF-CA3 synaptosomes from postnatal (P) day 28 hippocampal homogenate, a time-point at which MF-CA3 synapses have matured (Amaral and Dent, 1981). We used a previously established biochemical enrichment method (Taupin et al., 1994) that relies on the large size of MF-CA3 synapses (Figure S1A), and verified that this method enriches for MF-CA3 synaptosomes. Electron microscopy (EM) analysis showed the presence of large synaptosomes with a presynaptic compartment packed with synaptic vesicles engulfing a postsynaptic compartment (Figure S1B). To confirm the identity of these synaptosomes, we used the MF-CA3 synapse markers Synaptoporin (Synpr), a presynaptic vesicle-associated protein (Williams et al., 2011), and Nectin 3, a CA3-enriched (http://mouse.brain-map.org/; https://hipposeq.janelia.org/) (Cembrowski et al., 2016) adhesion protein that localizes to the postsynaptic side of PA junctions in mature MF-CA3 synapses (Mizoguchi et al., 2002) (Figure 1B). As an additional MF-CA3 synapse marker, we used glutamate receptor ionotropic kainate 5 (GluK5), a predominantly postsynaptically localized kainate receptor enriched at MF-CA3 synapses (Darstein et al., 2003; Petralia et al., 1994) (Figure S1C). Immunolabeling revealed the presence of all three MF-CA3 synaptic markers in synaptosomes (Figure S1D). Synpr labeling overlapped with the excitatory presynaptic marker vesicular glutamate transporter 1 (VGluT1), whereas Nectin 3 labeling was restricted to small puncta (Figure S1D), as expected for a PA-localized postsynaptic protein. GluK5 labeling was similarly restricted to small puncta (Figure S1D). Quantification showed an enrichment of large Synpr- and Nectin 3-positive synaptosomes in the MF-CA3 synaptosome-containing fraction (Figure S1E). We subsequently accelerated the biochemical enrichment procedure by omitting gradient centrifugation and depleting myelin from the sample (Figure S1F). Second, we subjected the biochemical preparation to fluorescence-activated synaptosome sorting (Biesemann et al., 2014) to further enrich for MF-CA3 synaptic material. To this end, we live-labeled MF-CA3 synaptosomes with a fluorophore-conjugated monoclonal antibody against the extracellular domain of Nectin 3 (Figure S1G) and with FM4-64 membrane dye to label plasma membrane-bound particles (Figure 1C; Figure S1H and I), and sorted them in a fluorescent cell sorter.

WB analysis showed the presence of MF-CA3 synapse markers Synpr and Nectin 3 in the sorted MF-CA3 synaptosome sample, whereas myelin and nuclear markers were largely depleted from the sorted fraction (Figure 1D). Immunofluorescence analysis confirmed the presence of Synpr-, Nectin 3-, and VGluT1-positive large synaptosomes in the sorted sample (Figure 1E and F). In addition to Synpr-/Nectin 3-/VGluT1-positive synaptosomes, VGluT1-positive particles were invariably present (Figure 1E and F). Such debris has previously been observed following fluorescent synaptosome sorting (Biesemann et al., 2014) and may represent synaptic membrane fragments resulting from the synaptosome preparation or sorting procedure, indicating that careful validation of potential hits resulting from proteomic analysis of this material is required. Together, these results show that a combination of biochemical enrichment, fluorescent labeling and sorting successfully enriches for MF-CA3 synaptosomes from hippocampal tissue homogenate.

### Proteomic analysis of isolated MF-CA3 synaptosomes

For proteomic analysis, we compared sorted MF-CA3 synaptosomes to P2 synaptosomes isolated in parallel from the same hippocampal homogenate (Figure S1F). P2 synaptosomes represent a mixed population of small hippocampal synapses and serve as a background reference in our analysis. We analyzed sorted MF-CA3 synaptosomes and P2 synaptosomes from three independent experiments (10-12 mice per experiment) by liquid chromatography tandem mass spectrometry (LC-MS/MS)-based proteomics. We identified 3592 proteins with at least 3 peptide identifications among replicates; 11,6% and 7,9% of these proteins were exclusively detected in sorted MF-CA3 synaptosomes or P2 synaptosomes, respectively (Figure 1G and Table S1). Gene ontology (GO) analysis showed a similar enrichment for synaptic terms in sorted MF-CA3 and P2 synaptosomes (Figure S1J). We calculated log_2_ fold-change (FC) enrichment in sorted MF-CA3 versus P2 synaptosomes using a label-free semi-quantitative approach based on the normalized spectral abundance factor (NSAF) (Figure 1H; Figure S1K and Table S1). Our analysis revealed 605 proteins significantly enriched in sorted MF-CA3 synaptosomes and 138 additional significant proteins exclusively detected in MF-CA3 synaptosomes (Figure 1H; Table S1). This collection comprises multiple proteins previously reported to be strongly enriched at MF-CA3 synapses, including the synaptic vesicle-associated proteins Synpr, Synapsin 3 (Syn3) (https://www.sysy.com), Rabphilin-3A (Rph3a) (Schlüter et al., 1999), and zinc transporter 3 (ZnT3/Slc30a3) (Wenzel et al., 1997); the glutamate receptors GluK2/Grik2 (Straub et al., 2016) and metabotropic glutamate receptor 3 (mGluR3/Grm3) (Shigemoto et al., 1997); the presynaptic scaffold protein liprin-α2 (Ppfia2) (Spangler et al., 2011); the dense core vesicle secretion-related protein calcium-dependent secretion activator 2 (CAPS2/Cadps2) (Sadakata et al., 2006); and the PA-associated proteins Nectin 3 (listed as Pvrl3 in Table S1) and afadin (Af-6/Mllt4) (Mizoguchi et al., 2002) (Figure 1H; Figure S1K). Relatively few peptides were detected for Nectin 3, the marker used to label MF-CA3 synaptosomes, which may be due to a low abundance and restricted localization of this protein (Figure S1D). Supporting this notion, similar peptide amounts were detected for GluK5/Grik5 (Table S1), a transmembrane receptor with a similarly restricted localization in MF-CA3 synaptosomes (Figure S1D). Using immunohistochemistry (IHC), we confirmed strongly enriched localization of Syn3, Af-6, mGluR2/3, and ZnT3 in SL (Figure 1I). Together, these results indicate that LC-MS/MS analysis of biochemically enriched and fluorescently sorted MF-CA3 synaptosomes confidently identifies MF-CA3 synaptic proteins.

### Characterization of MF-CA3 synaptosome CSP composition

Using the UniProt database (https://www.uniprot.org), we next queried our proteomic dataset for transmembrane, membrane-anchored, and secreted proteins among the proteins significantly enriched in sorted MF-CA3 synaptosomes and those exclusively detected in MF-CA3 synaptosomes. We identified a panel of 77 CSPs including adhesion proteins, receptors, secreted glycoproteins, receptor protein tyrosine phosphatases and tyrosine kinases (Figure 2A and Table S2). Most major CSP families, such as the immunoglobulin (Ig) superfamily (IgSF), fibronectin type-III (FN3), and leucine-rich repeat (LRR) family, were represented (Figure 2B). Only a small proportion (20,8%) of these MF-CA3 synapse CSP candidates has been previously reported to localize or function at MF-CA3 synapses (Table S2). Using the synapse biology GO database SynGO (https://syngoportal.org) (Koopmans et al., 2019), we determined that 52% of the identified CSP genes has a SynGO annotation (Figure 2C), indicating that these are known synaptic CSPs. The secreted protein bone morphogenic protein/retinoic acid-inducible neural-specific protein 2 (BRINP2), and the transmembrane receptors family with sequence similarity 171 member A2 (FAM171A2), adipocyte plasma membrane-associated protein (APMAP), and immunoglobulin superfamily 8 (IgSF8) lack a function in the brain altogether (Figure 2C).

**Figure 2.**
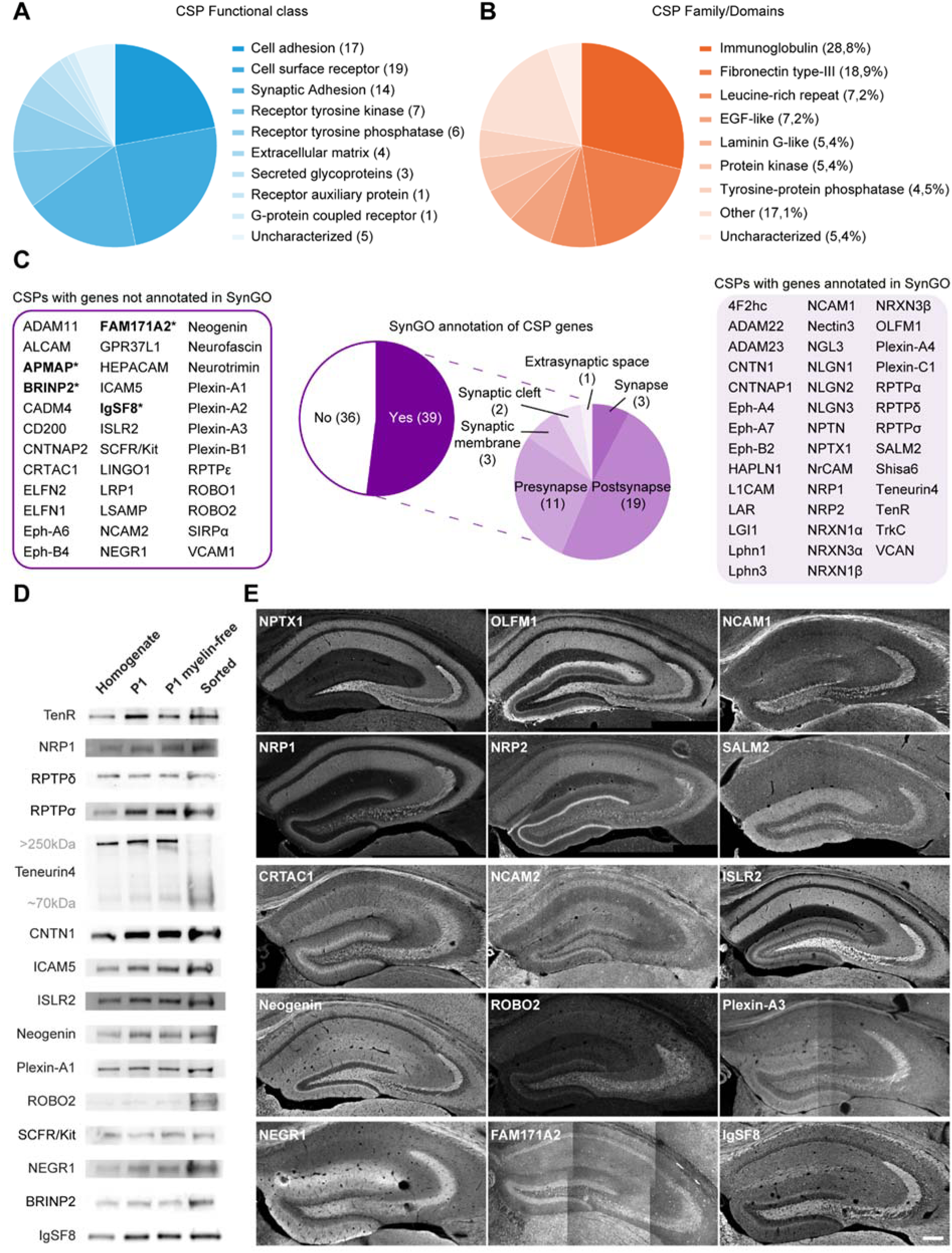
Characterization of MF-CA3 synaptosome CSP composition. (**A**) Protein functional classes represented in the group of CSPs identified in isolated MF-CA3 synaptosomes. (**B**) Occurrence of protein domains among identified CSPs. (**C**) SynGO cellular component analysis of CSP genes (75) and respective CSPs (77). Discrepancy between gene number and CSP number is related to NRXN isoforms. Proteins with unknown function in the brain are highlighted in bold with an asterisk: APMAP, BRINP2, FAM171A2 and IgSF8. (**D**) Validation of 15 CSPs in sorted MF-CA3 synaptosomes by western blot including BRINP2 and IgSF8, CSPs without a known function in the brain. (**E**) Confocal images of P28 mouse hippocampal sections immunostained for 15 CSPs showing a striking laminar distribution to SL. Of the CSPs without a known function in the brain, IgSF8 showed strong labeling in the MF-CA3 pathway. Scale bar in (E) 200 μm. See also Figure S2.

To validate these results, we tested a large panel of antibodies for detection of CSPs by WB and IHC (Figure S2A, Table S2). We validated the presence of 15 CSPs in sorted P28 MF-CA3 synaptosomes by WB (Figure 2D). Of the SynGO-annotated CSPs, we confirmed the secreted ECM protein TenR; the transmembrane receptors neuropilin-1 (NRP1), receptor-type tyrosine-protein phosphatase delta (RPTPδ) and sigma (RPTPσ), and Teneurin 4; as well as the GPI-anchored receptor contactin-1 (CNTN1) (Figure 2D). Of the CSPs lacking a SynGO-annotated synaptic function, we validated the transmembrane receptors intercellular adhesion molecule 5 (ICAM5), immunoglobulin superfamily containing leucine-rich repeat protein 2 (ISLR2), Neogenin, Plexin-A1, roundabout homolog 2 (ROBO2), mast/stem cell growth factor receptor SCFR/Kit; and the GPI-anchored receptor neural growth regulator 1 (NEGR1) (Figure 2D). Of the CSPs without function in the brain, we confirmed the presence of BRINP2 and IgSF8 in sorted MF-CA3 synaptosomes (Figure 2D).

Using IHC on P28 mouse hippocampal sections, we validated localization to the MF-CA3 pathway for 21 CSPs, 15 of which showed a striking laminar distribution (Figure 2E; Figure S2B). We confirmed strong localization to SL for the SynGO-annotated secreted proteins neuronal pentraxin-1 (NPTX1) and neuronal olfactomedin-related ER localized protein/olfactomedin-1 (OLFM1); as well as the transmembrane receptors neural cell adhesion molecule 1 (NCAM1), NRP1 and NRP2, and leucine-rich repeat and fibronectin type III domain-containing protein 1 (LRFN1/SALM2) (Figure 2E). Of the CSPs lacking a SynGO-annotated synaptic function, the secreted protein cartilage acidic protein 1 (CRTAC1); the transmembrane receptors NCAM2, ISLR2, Neogenin, ROBO2, and Plexin-A3; and the GPI-anchored receptor NEGR1 displayed strongly enriched immunoreactivity in SL (Figure 2E). Of the CSPs without a known function in the brain, we validated FAM171A2 in SL, which displayed weak labeling in SL, and IgSF8, which showed strong labeling in the MF-CA3 pathway (Figure 2E).

Besides these 15 CSPs with lamina-specific localization patterns, we observed a broad hippocampal distribution, including SL, for an additional 6 CSPs (Figure S2B) that were identified in sorted MF-CA3 synaptosomes (Figure 2C; Table S1). Among these 6 CSPs are CNTN1 and TenR, which we also confirmed by WB (Figure 2D). CSPs detected in sorted MF-CA3 synaptosomes by WB and localized to SL by IHC were in good agreement (Figure 2D and E; Figure S2B), with three exceptions: ICAM5, Plexin-A1, and SCFR/Kit. While these 3 CSPs were validated in sorted MF-CA3 synaptosomes by WB (Figure 2D), they did not show immunoreactivity in SL (Figure S2C). It is possible that the antibodies used do not recognize the epitopes for these CSPs in SL or that these CSPs are present in MF-CA3 sorted synaptosomes as contaminants and represent false positives. SCFR/Kit however has previously been shown to function at MF-CA3 synapses (Katafuchi et al., 2000; Kondo et al., 2002; Motro et al., 1996) (Table S2). In conclusion, validation of the MF-CA3 synaptosome proteome analysis confirms the presence of a diverse CSP repertoire at mature MF-CA3 synapses, including several that lack a function in the brain (BRINP2, FAM171A2, IgSF8).

### Interactome analysis of CSPs identified in sorted MF-CA3 synaptosomes

CSPs exert their function by forming protein-protein interaction networks with other CSPs (Südhof, 2018; de Wit and Ghosh, 2016). To elucidate ligand-receptor relationships among the CSPs identified in sorted MF-CA3 synaptosomes and obtain clues about potential roles of the CSPs with unknown brain function, we systematically screened for pairwise interactions. We generated a library of constructs containing the extracellular domains of 73 CSPs (see Methods) identified in sorted MF-CA3 synaptosomes (Figure 2C; Table S2), fused at their C-terminus with human alkaline phosphatase (AP) or the Fc region of human IgG1. We subsequently generated and validated recombinant AP- and Fc-tagged proteins (Table S3 and Methods), and performed a pairwise high-throughput interaction screen using a semi-automated 384-well format enzyme-linked immunosorbent assay (ELISA)-based assay (Ozgul et al., 2019; Özkan et al., 2013; Wojtowicz et al., 2007) (Figure 3A and B). The screen was designed to test interactions between 73 AP- and Fc-tagged proteins in both orientations (i.e. AP-X vs Fc-Y and AP-Y vs Fc-X), resulting in 73 x 73 = 5,329 experimental points and screening of 2701 unique pairwise interactions. Positive interactions (blue in Figure 3C) were defined as those wells that exhibited an OD_650_ >5-fold over background values (FOB) and that were not present in every bait. The vast majority were negative wells, which were either completely clear, or had negligible FOB values (white in Figure 3C). False-positive wells were characterized by having significant signal in all baits (not shown). Combining the results of three independent experiments, using the criteria outlined above and considering only interactions detected at least twice independent of orientation, we identified 38 interacting pairs, 10 of which, to the best of our knowledge, have not been reported before (Figure 3C; Table S3).

**Figure 3.**
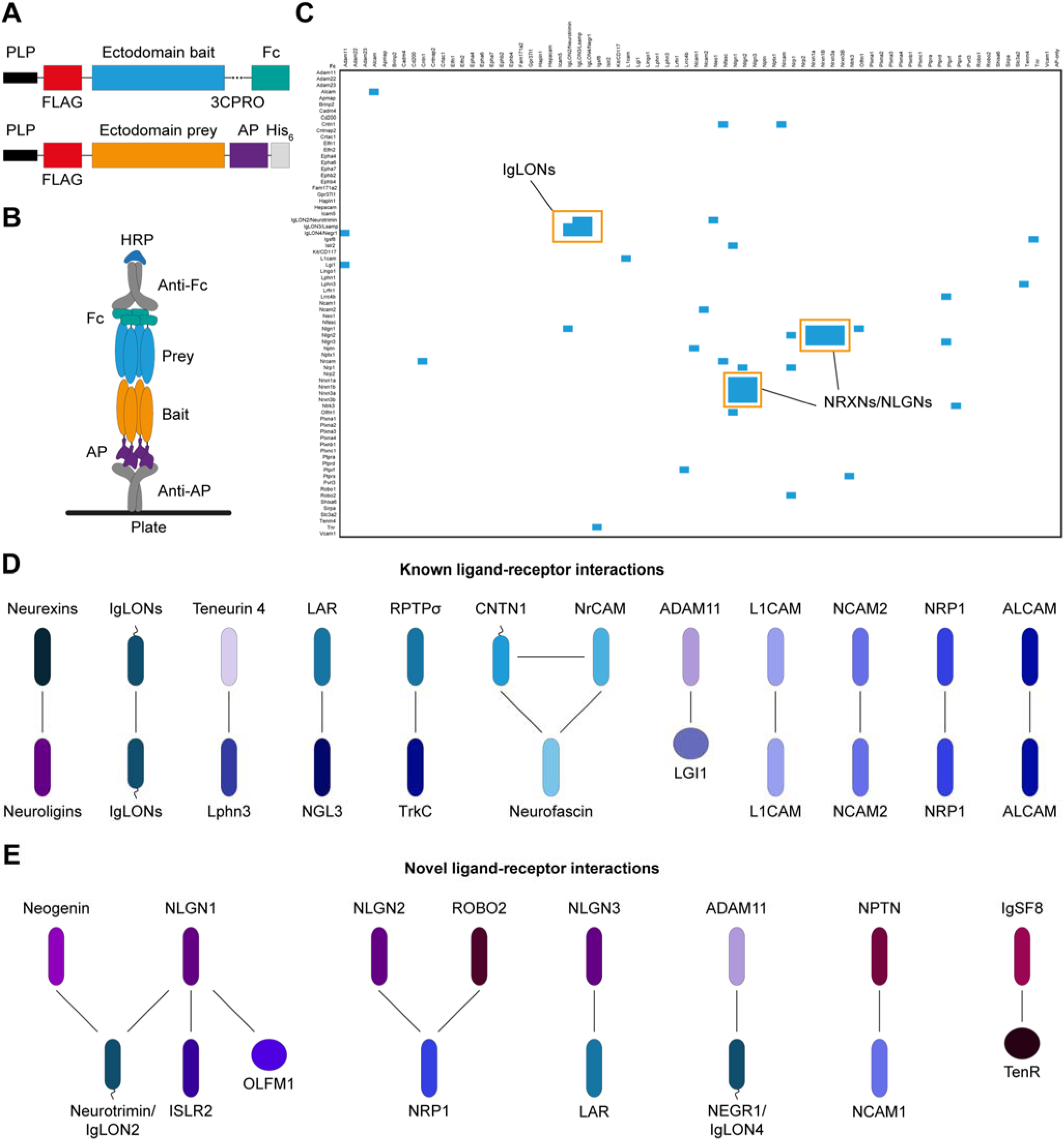
Interactome analysis of CSPs identified in sorted MF-CA3 synaptosomes. (**A**) Schematic diagram of the constructs used to perform the CSP interactome analysis. Ectodomains of transmembrane and GPI-anchored proteins, or full-length protein in case of secreted proteins, were C-terminally tagged with human Fc fragment or alkaline phosphatase (AP). Both constructs have a leader peptide (PLP) to ensure that recombinant proteins are secreted. The Fc-fusion has a 3CPro cleavage site for purification purposes, whereas the AP-fusion construct has an additional His tag. (**B**) Schematic diagram of the ELISA-based assay showing the orientation of bait and prey recombinant proteins. (**C**) Matrix of the data obtained from interactions between 73 AP- and Fc-tagged proteins in both orientations (i.e. AP-X vs Fc-Y and AP-Y vs Fc-X), resulting in 73 x 73 = 5,329 experimental points and 2701 unique pairwise interactions. Columns contain AP-fusion baits, including the AP-only construct as negative control, whereas rows contain Fc-fusion preys. Positive ligand-receptor interactions are indicated in blue. Two major modules of known ligand-receptor pairs are highlighted in orange: the NRXNs-NLGNs and the IgLON family. (**D**) and (**E**) Interaction networks of known and novel ligand-receptor pairs identified, respectively. Among novel receptor-ligand pairs identified, the interaction between IgSF8 and TenR was the only one including a CSP of uncharacterized brain function.

The ELISA-based interaction screen reproducibly identified two major modules of known ligand-receptor pairs: the neurexins and neuroligins (Südhof, 2017), and the IgLON family (Ranaivoson et al., 2019), confirming specificity of the screen. Multiple additional known interactions were reproducibly identified, including those of Teneurin 4 and latrophilin 3 (Lphn3) (Boucard et al., 2014; O’Sullivan et al., 2012; Silva et al., 2011), and of leukocyte common antigen-related (LAR) and netrin-G ligand 3 (NGL3) (Woo et al., 2009) (Figure 3C and D; Table S3). Of the novel receptor-ligand pairs we identified, the interaction between IgSF8 and TenR was among the most reproducible (Figure 3C and E; Table S3). Of the MF-CA3 synaptosome CSPs wihout a brain function, IgSF8 was the only CSP for which the interactome screen yielded a binding partner (Figure 3C and E). Taken together, the CSP interactome analysis identifies multiple ligand-receptor pair modules at MF-CA3 synapses, including several novel ones, that may play a role in shaping MF-CA3 synapse structure and function.

### The secreted ECM protein TenR is a novel ligand for IgSF8

The results from our interactome screen identified TenR as a novel ligand for the uncharacterized neuronal receptor IgSF8. As this screen encompassed only those CSPs that reached significance in our proteomic analysis, we next performed affinity chromatography using recombinant IgSF8-ecto-Fc as bait on whole brain synaptosome extract as well as MF-CA3 synaptosome extract to screen a larger panel of prey proteins (Figure 4A and Table S4). Bound proteins were analyzed by LC-MS/MS (Savas et al., 2014). We identified TenR as the main CSP interacting with IgSF8, both in whole brain synaptosome extract, as well as in MF-CA3 synaptosome extract (Figure 4B and C; Table S4).

**Figure 4.**
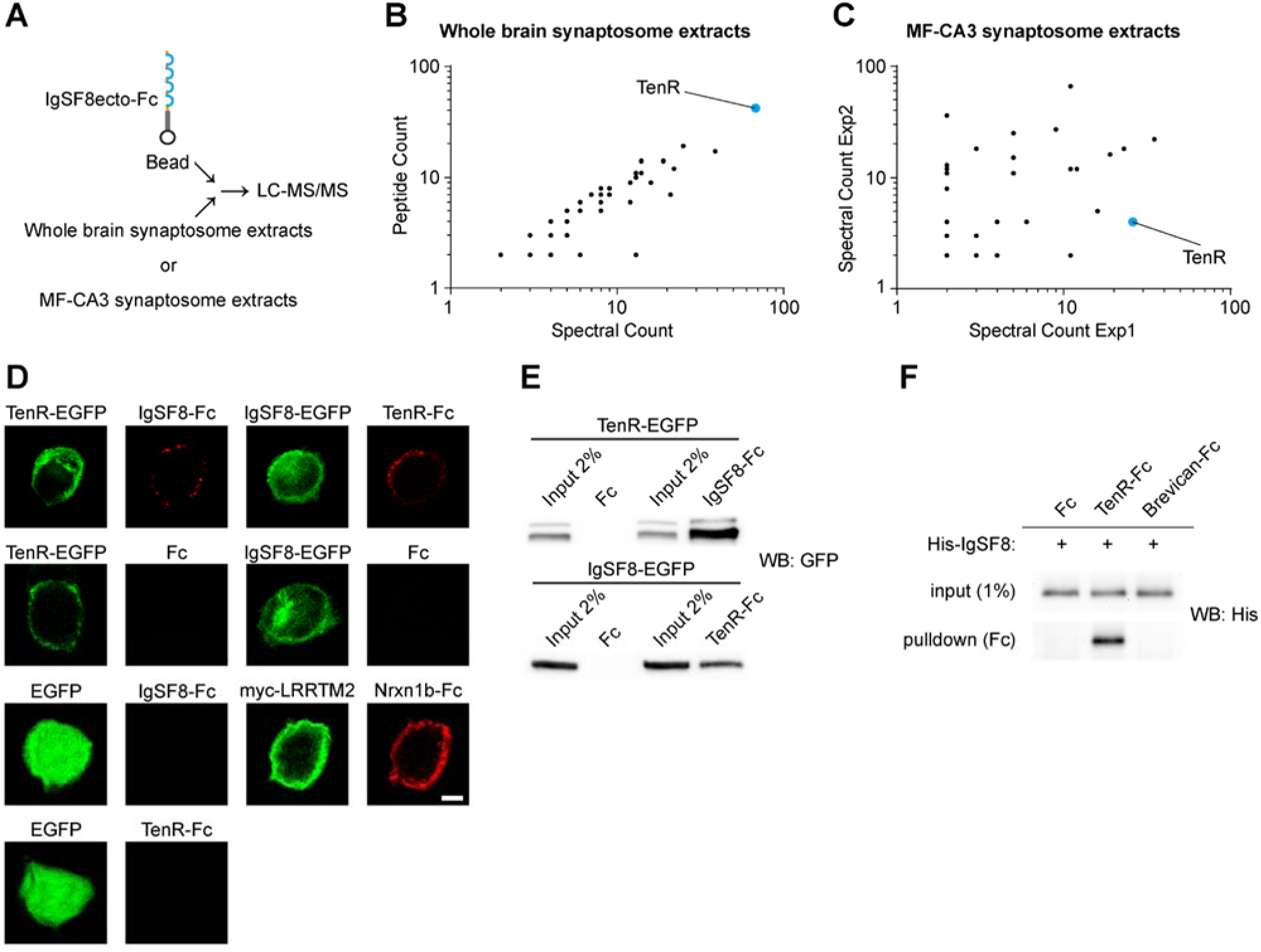
The secreted ECM protein TenR is a novel ligand for IgSF8. (**A**) Cartoon illustrating the Ecto-Fc MS workflow. (**B**) Graph of spectral and peptide counts of proteins captured in one independent pull-down experiment using whole rat brain synaptosome lysates. Only proteins which were absent in Fc controls and present with ≥ 2 spectral counts per IgSF8-ecto-Fc experiment are included. TenR is highlighted in blue. (**C**) Graph of spectral counts of proteins captured in two independent pull-down experiments using P1 MF-CA3 synaptosome lysates. Only proteins with ≤ 1 spectral counts in Fc controls and ≥ 2 spectral counts per IgSF8-ecto-Fc experiment are included. TenR is highlighted in blue. (**D**) Confocal images of cell-surface binding assays in transfected HEK293T cells show IgSF8-TenR interaction. Negative and positive controls used in the cell-surface binding assays are included. (**E**) Pull-down assays in transfected HEK293T cells show IgSF8-TenR interaction. (**F**) Direct binding assays show direct and specific interaction of IgSF8 with TenR but not with ECM protein Brevican. Scale bar in (D) 5 μm.

To validate the IgSF8-TenR interaction, we used cell-surface binding assays and pull-down assays in transfected HEK293T cells and confirmed that IgSF8 and TenR bind to each other (Figure 4D and E). To test whether the IgSF8-TenR interaction is direct, we mixed equimolar amounts of Fc control protein, TenR-Fc, or the ECM protein Brevican-Fc, with His-IgSF8 recombinant protein and precipitated Fc proteins. This binding assay showed that the IgSF8-TenR interaction is direct and specific (Figure 4F). Taken together, the results from CSP interactome screening, affinity chromatography coupled with LC-MS/MS analysis, and binding assays using HEK293T cells or recombinant proteins identify IgSF8-TenR as a novel receptor-ligand pair at MF-CA3 synapses.

### Loss of IgSF8 impairs MF-CA3 synapse architecture and filopodia density

Proteomic analysis of sorted MF-CA3 synaptosomes, validation experiments, and CSP interactome screening identified IgSF8 as a receptor of unknown function enriched at MF-CA3 synapses and the secreted ECM protein TenR as a novel IgSF8 ligand. The identification of TenR, which regulates synaptic structure, plasticity, and excitability (Brenneke et al., 2004; Nikonenko et al., 2003; Saghatelyan et al., 2001), as a ligand for IgSF8 suggests that IgSF8 may function in synapse development. IgSF8 is a type I membrane protein with a large extracellular domain containing four Ig-like C2-type domains, a transmembrane region, and a short cytoplasmic domain (Figure 5A). This protein is one of four members of an IgSF subfamily containing the Glu-Trp-Ile (EWI) transmembrane motif of unknown biological function.

**Figure 5.**
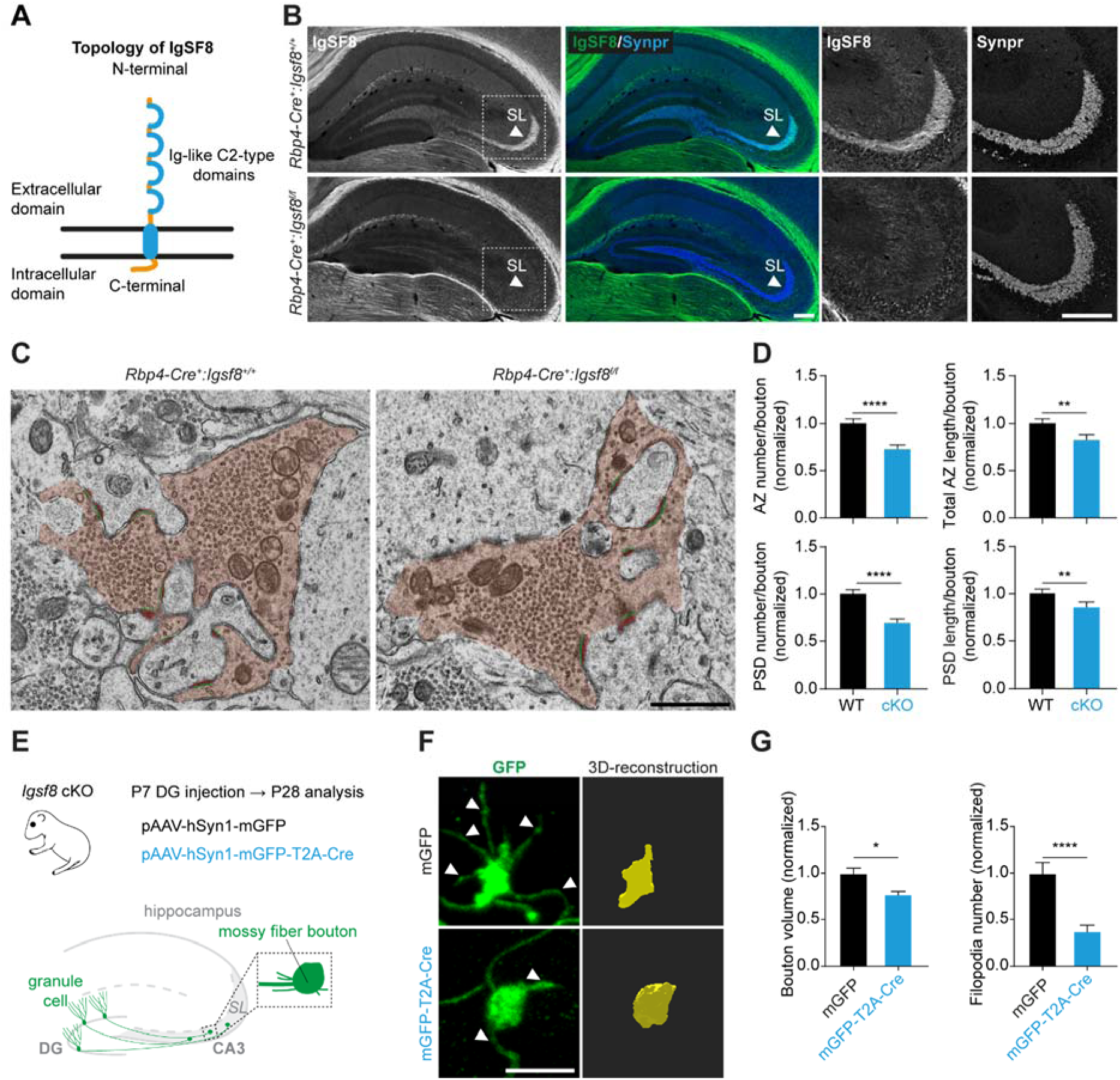
Loss of IgSF8 impairs MF-CA3 synapse architecture and filopodia density. (**A**) Cartoon of IgSF8 protein topology. (**B**) Confocal images of P28 *Rbp4-Cre*:*Igsf8* cKO mouse hippocampal sections immunostained for IgSF8 and Synpr. Arrowheads indicate SL. Magnified insets of the SL in CA3 are shown on the right. (**C**) Electron microscope images of MF-CA3 synapses from *Rbp4-Cre*:*Igsf8* cKO and WT littermates (5000x magnification). MF boutons are highlighted in orange. AZs and PSDs are highlighted in green and red, respectively. (**D**) Graphs show quantification of analysis done in (C) using three littermate mice per condition (WT, n = 135 boutons; cKO, n = 136). (**E**) Experimental design to analyze structural changes in MF boutons following deletion of *Igsf8* specifically in DG granule cells. (**F**) Stacks of confocal images of individual MF boutons (left) and respective 3D reconstructions (right) to analyze MF bouton volume, surface area and number of filopodia. Arrowheads show filopodia emerging from MF boutons. (**G**) Graphs show quantification of analysis done in (F) using three littermate mice per condition (mGFP, n=27 boutons; mGFP-T2A-Cre n=29). Graphs show mean ± SEM. Mann-Whitney tests were used; *\*P* < 0.05; *\*\*P* < 0.01; *\*\*\*\*P* < 0.0001. Scale bars in (B) 200 μm, in (C) 1 μm, and in (F) 5 μm. See also Figure S3.

To start elucidating the role of IgSF8 at MF-CA3 synapses, we first assessed its protein expression profile and localization. We analyzed IgSF8 expression in hippocampal lysates and found that IgSF8 protein levels mildly increased during postnatal development (Figure S5A). To determine the synaptic localization of IgSF8, we performed subcellular fractionation, as the conditions required for IgSF8 IHC proved unsuitable for high-resolution imaging. IgSF8 mainly distributed to the Triton-soluble fraction of synaptosomes containing the presynaptic protein synaptophysin (Figure S5B). In addition, two faint bands were observed in the Triton-insoluble fraction containing the postsynaptic protein PSD95 (Figure S5B). These biochemical results support a predominantly presynaptic localization of IgSF8, but do not exclude a postsynaptic presence.

To determine the functional significance of IgSF8 for MF-CA3 synapse development, we removed IgSF8 presynaptically by crossing *Igsf8* conditional knockout (cKO) mice (Inoue et al., 2012) with the dentate GC-specific *Rbp4*-*Cre* line. This largely and selectively abolished IgSF8 immunoreactivity in SL, in agreement with a predominantly presynaptic localization of IgSF8 at MF-CA3 synapses. *Igsf8* cKO did not affect gross MF-CA3 tract morphology (Figure 5B). Infecting cultured *Igsf8* cKO hippocampal neurons with a lentiviral (LV) vector harboring Cre recombinase strongly reduced IgSF8 protein levels (Figure S5C), further validating the utility of the *Igsf8* cKO line for assessing IgSF8 function.

To analyze MF-CA3 synaptic architecture, we prepared hippocampal sections from P30 *Rbp4-Cre:Igsf8* cKO and control littermates and imaged synapses by transmission electron microscopy (TEM) (Figure 5C). MF-CA3 bouton number and area were not affected in *Igsf8* cKO mice (Figure S5D and E). We observed a clear reduction in active zone (AZ) number and length, and a corresponding decrease in postsynaptic density (PSD) number and length (Figure 5D), indicating that loss of IgSF8 impairs MF-CA3 synaptic architecture.

To analyze MF bouton morphology, we injected adeno-associated viral (AAV) vectors expressing membrane-GFP (mGFP) and Cre recombinase in P7 *Igsf8* cKO mice to remove IgSF8 expression and used AAV-mGFP as a control (Figure 5E). We imaged P28 MF boutons using confocal microscopy and reconstructed them in 3D. We observed a mild reduction in volume of the main MF bouton (Figure 5F and G). In addition, we observed a dramatic decrease in the number of filopodia emerging from the main bouton in *Igsf8* cKO mice compared to controls (Figure 5F and G). Together, these results indicate that IgSF8 plays a role in shaping MF-CA3 synapse architecture, bouton morphology and filopodia density.

### Decreased spontaneous transmission in Igsf8 cKO CA3 neurons

To analyze functional changes at MF-CA3 synapses in the absence of IgSF8, we performed whole-cell voltage-clamp recordings of CA3 neurons in acute hippocampal slices from P27-35 *Rbp4-Cre:Igsf8* cKO and control littermates. Frequency and amplitude of spontaneous excitatory postsynaptic currents (sEPSCs) were reduced in *Igsf8* cKO mice compared to controls (Figure 6A-E), whereas sEPSC decay time was not affected (Figure S6A). Histogram analysis showed a loss of large-amplitude sEPSCs in *Igsf8* cKO mice, (Figure S6B and C), suggesting an impairment of MF-CA3 synapses that give rise to these events (Henze et al., 2002). These observations are consistent with the decrease in number and length of MF-CA3 synaptic junctions we observed in *Igsf8* cKO mice.

**Figure 6.**
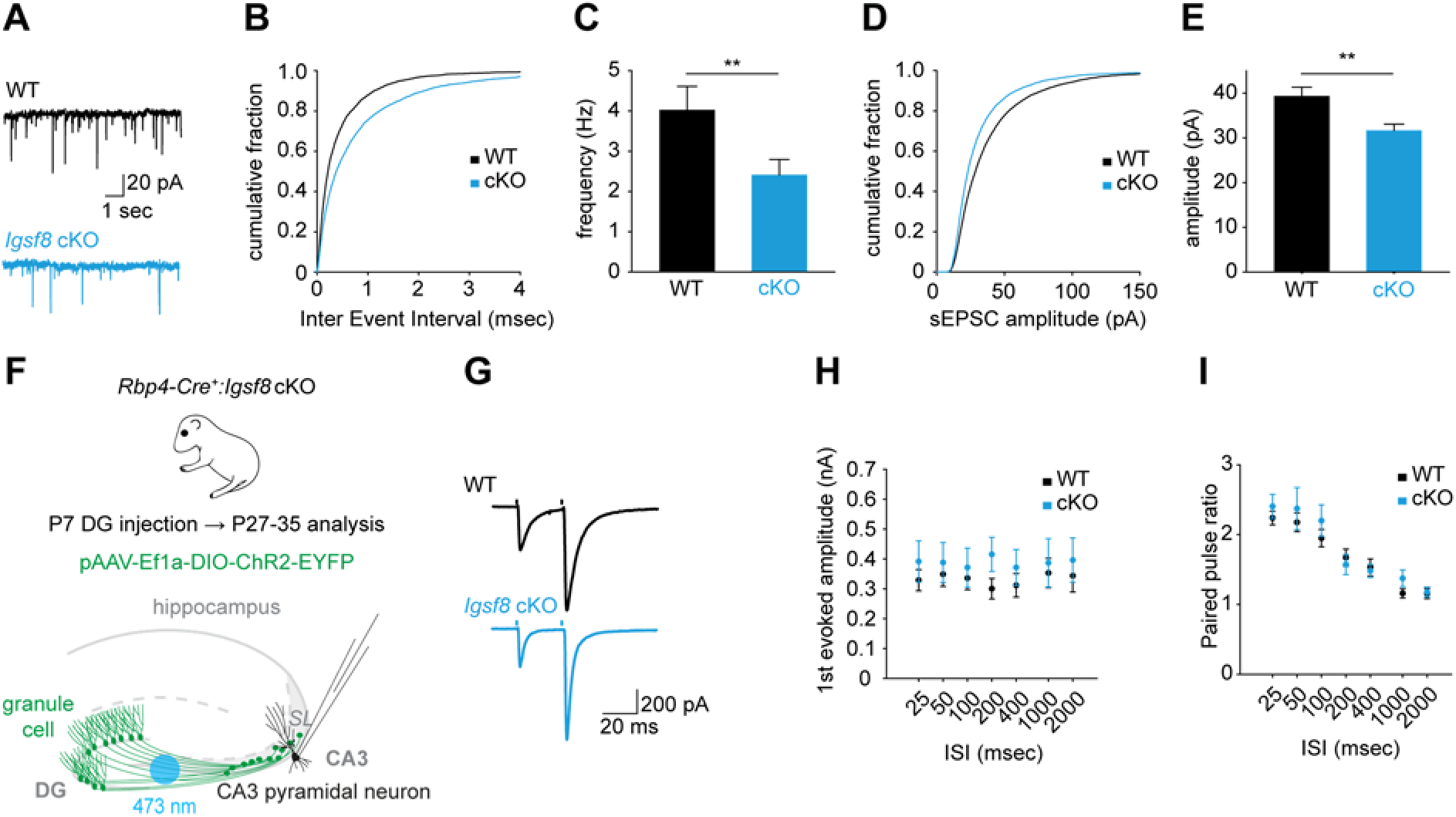
Decreased spontaneous transmission in *Igsf8* cKO CA3 neurons. (**A**) Representative sEPSC traces from whole-cell voltage-clamp recordings of CA3 neurons in acute hippocampal slices of *Rbp4-Cre*:*Igsf8* cKO and WT littermates. (**B**) Cumulative distribution of sEPSC inter-event intervals. (**C**) Quantification of sEPSC frequency. (**D**) Cumulative distribution of sEPSC amplitudes. (**E**) Quantification of sEPSC amplitudes. (**F**) Cartoon illustrating whole-cell voltage-clamp recordings of CA3 neurons to measure MF-evoked responses in acute hippocampal slices of *Rbp4-Cre*:*Igsf8* cKO and WT littermates using optogenetics. (**G**) Representative traces of paired-pulse ratio in *Rbp4-Cre*:*Igsf8* cKO and WT littermates using optogenetics. (**H**) Quantification of first evoked amplitudes. (**I**) Quantification of paired-pulse ratios. Three littermate mice were used per condition. For sEPSCs: WT, n = 31 neurons and cKO, n = 38. For paired-pulse ratio: WT, n = 23 neurons and cKO, n= 21. Graphs show mean ± SEM. Mann-Whitney tests were used in (C) and (E). *\*\*P* < 0.01. See also Figure S4.

To specifically assess synaptic transmission at MF-CA3 synapses, we expressed Cre-dependent Channelrhodopsin-2 (DIO-ChR2) in *Rbp4-Cre:Igsf8* cKO and control littermates and optically stimulated MF axons while performing whole-cell voltage-clamp recordings from CA3 pyramidal neurons (Figure 6F). Using this approach, only MFs of GCs lacking IgSF8 are stimulated. We analyzed paired-pulse facilitation, a form of short-term plasticity, to assess MF-CA3 presynaptic properties, but found no differences in the amplitude of the first evoked EPSC or paired-pulse ratio between *Igsf8* cKO and control mice (Figure 6G-I). Thus, our results show that loss of presynaptic IgSF8 impairs spontaneous synaptic transmission but does not alter evoked transmission at MF-CA3 synapses.

### Igsf8 cKO reduces feedforward inhibition and increases excitability of CA3 neurons

The filopodia emanating from MF boutons synapse onto interneurons in SL that mediate feedforward inhibition of CA3 neurons (Acsády et al., 1998; Rebola et al., 2017; Torborg et al., 2010). Given the robust decrease in the number of filopodia in boutons of *Igsf8* cKO mice (Figure 5F and G), we hypothesized that stimulation of *Igsf8* cKO MFs would result in reduced recruitment of feedforward inhibition onto CA3 neurons. To test this, we recorded light-evoked EPSCs and IPSCs from the same CA3 pyramidal neuron in *Rbp4-Cre:Igsf8* cKO and control littermates. Consistent with our hypothesis, we observed a robust increase in the excitation-inhibition ratio of CA3 pyramidal neurons in mice lacking presynaptic IgSF8 compared to control mice (Figure 7A-D).

**Figure 7.**
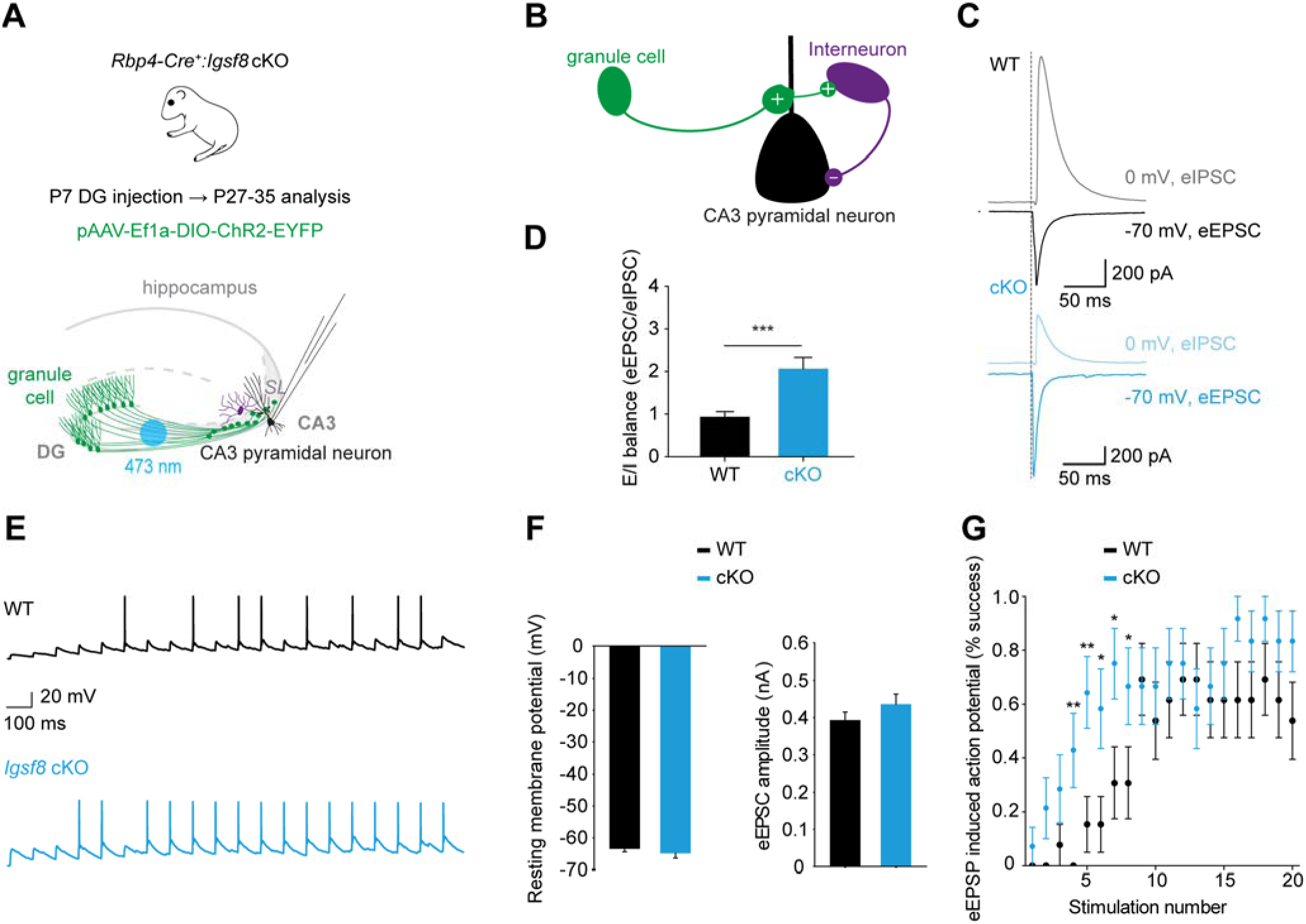
*Igsf8* cKO reduces feedforward inhibition and increases excitability of CA3 neurons. (**A**) Cartoon illustrating whole-cell voltage-clamp recordings of CA3 neurons to measure MF-evoked responses in acute hippocampal slices of *Rbp4-Cre*:*Igsf8* cKO and WT littermates using optogenetics. (**B**) Cartoon illustrating feedforward inhibition microcircuit in CA3. Plus and minus signs represent excitatory and inhibitory synapses, respectively. (**C**) Representative eEPSC and eIPSC traces from CA3 pyramidal neurons in WT or cKO mice. (**D**) Quantification of excitation-inhibition balance in CA3 neurons in WT and cKO mice. (**E**) Representative eEPSP traces from CA3 pyramidal neurons in WT or cKO mice. (**F**) Quantification of resting membrane potentials and eEPSC amplitudes of CA3 neurons in WT and cKO mice. (**G**) Quantification of induced action potential firing in CA3 neurons in WT and cKO mice after a 10 Hz train of 20 stimuli. Three littermate mice were used per condition. For eEPSCs and eIPSCs: WT, n = 35 neurons and cKO, n= 29. For eEPSPs: WT, n = 13 neurons and cKO, n= 14. Graphs show mean ± SEM. Mann-Whitney tests were used in (D). Student’s t-test was used in (G). *\*P* < 0.05; *\*\*P* < 0.01; *\*\*\*P* < 0.001.

Feedforward inhibition mediated by MF filopodia plays an important role in the control of CA3 pyramidal neuron excitability (Torborg et al., 2010). To determine the consequences of reduced feedforward inhibition in *Igsf8* cKO mice on CA3 pyramidal neuron excitability, we optically stimulated MF axons and analyzed action potential (AP) firing in CA3 neurons in *Rbp4-Cre:Igsf8* cKO and control littermates (Figure 7E). In this experiment, we first recorded CA3 neurons in voltage-clamp mode at −82 mV (the Cl^−^ reversal potential) and stimulated MFs to establish a comparable eEPSC amplitude of ±400 pA in both conditions (Figure 7F). CA3 neurons were then switched to current-clamp mode to record resting membrane potential, which was comparable between both groups (Figure 7G), followed by optical stimulation of MFs with a 10Hz train of 20 stimuli. We found that CA3 neurons in *Igsf8* cKO mice started firing APs earlier during the train than in control littermates (Figure 7G), supporting the notion that reduced feedforward inhibition in the absence of IgSF8 leads to increased MF-induced excitability of CA3 neurons. Taken together, these results show loss of IgSF8 reduces MF bouton filopodia density, resulting in a decrease in feedforward inhibition and an increase in excitability of CA3 pyramidal neurons, indicating that IgSF8 is an organizer of hippocampal CA3 microcircuit connectivity and function.

## Discussion

Synaptic diversity is a key organizational feature of neural circuits and plays a critical role in information processing in neurons. Its underlying molecular basis remains poorly understood. Here, we surveyed the proteome of isolated MF-CA3 synaptosomes to identify cell-surface regulators of MF-CA3 synapse properties. Our results reveal a diverse repertoire of CSPs at MF-CA3 synapses and identify the neuronal receptor IgSF8 as a novel regulator of CA3 microcircuit connectivity and function.

### Identification of a diverse CSP repertoire at MF-CA3 synapses

We combined biochemical enrichment, antibody labeling of a synapse type-enriched surface marker, and fluorescent sorting to isolate MF-CA3 synaptosomes. We reasoned that, despite relatively harsh synaptosome preparation and fluorescent sorting procedures, this approach coupled with highly sensitive MS analysis would allow us to gain insight into CSP composition of a specific excitatory synapse type and identify novel regulators of MF-CA3 synapse properties.

We identified a rich CSP repertoire at MF-CA3 synapses that includes adhesion proteins, guidance cue receptors, synaptic cleft and ECM proteins, as well as several CSPs of uncharacterized brain function. Biochemical validation experiments confirmed the presence of 15 CSPs in MF-CA3 synaptosomes. IHC experiments localized 21 CSPs to the MF-CA3 pathway. Most of these showed striking lamina-specific distributions and enriched localization to SL, whereas others displayed a broader laminar distribution. CSP interactome screening revealed multiple ligand-receptor modules among the MF-CA3 synaptosome CSPs, including several novel ones. The combined results of our proteomic analysis, validation experiments, and interactome screen suggest that a combinatorial code of CSP modules defines MF-CA3 synapse identity. This is reminiscent of the combinatorial molecular codes of broadly and discretely expressed CSPs that specify cell-type identity (Földy et al., 2016; Li et al., 2017; Shekhar et al., 2016).

Several CSPs with a role in MF-CA3 synapse development, including the secreted glycoprotein C1QL3 (Matsuda et al., 2016), Cadherin-9 (CDH9) (Williams et al., 2011), and the G-protein-coupled receptor-like protein GPR158 (Condomitti et al., 2018), were detected in sorted MF-CA3 synaptosomes (Table S1) but did not reach significance. This suggests that the cell-surface proteome of MF-CA3 synapses is likely more diverse than our results indicate. In part, this diversity may reflect heterogeneity within the MF-CA3 synapse population. The maturational state of MF-CA3 synapses varies due to the continuous integration of newborn GCs into the hippocampal circuit (Toni et al., 2008). Different histories of synaptic activity may also diversify CSP composition.

Our CSP interactome screen identified 10 novel interactions, 9 of which will need to be validated by independent methods. As the screen was restricted to those CSPs that reached significance in our proteome analysis, tests only binary interactions, does not take splice variants into account, and does not detect binding affinities weaker than ±10 μM (Ozgul et al., 2019; Özkan et al., 2013; Ranaivoson et al., 2019; Visser et al., 2015), the MF-CA3 synapse CSP interactome is likely more complex. To what extent developmental stage and neural activity influence cell-surface composition and interactome of MF-CA3 synapses will be a topic of future research.

### CSPs of uncharacterized function at MF-CA3 synapses

Our proteomic survey of isolated MF-CA3 synaptosomes uncovered several CSPs of uncharacterized function in the brain (APMAP, BRINP2, FAM171A2, IgSF8). The transmembrane protein APMAP interacts with the amyloid precursor protein (APP) and the γ-secretase complex and negatively regulates amyloid-β production (Gerber et al., 2019; Mosser et al., 2015). *Apmap* KO mice display spatial memory defects (Gerber et al., 2019). The transmembrane protein FAM171A2 induces membrane protrusions in cultured cells upon overexpression and is required for invasive growth of melanoma cells (Rasila et al., 2019). FAM171A2 was identified in the excitatory synaptic cleft proteome (Loh et al., 2016) and localizes to SL, but its role there remains unknown. BRINP2 is a secreted glycoprotein that is highly expressed in CA3 (Kawano et al., 2004) and detected in MF-CA3 synaptosomes. *Brinp2* KO mice display hyperactivity (Berkowicz et al., 2016). Multiple secreted and ECM proteins were validated at MF-CA3 synapses (BRINP2, CRTAC1, NPTX1, OLFM1 and TenR), of which the synaptic function of CRTAC1 is not known. CRTAC1 acts as a Nogo receptor-1 antagonist in regulating lateral olfactory axon tract bundling in early development (Sato et al., 2011). Interestingly, multiple CSPs with roles in early stages of circuit development (NRP1, NRP2, CRTAC1, ISLR2, Neogenin, ROBO2, Plexin-A1, and Plexin-A3) were validated at mature MF-CA3 synapses. With the exception of the neuropilins (Demyanenko et al., 2014; Morita et al., 2006; Tran et al., 2009), their synaptic role remains unknown.

### Identification of IgSF8 as a regulator of CA3 microcircuit development

The combined results of our proteomics analysis, validation experiments, and interactome screen identified IgSF8, a member of the enigmatic Glu-Trp-Ile (EWI) IgSF subfamily, as an uncharacterized neuronal receptor that is highly enriched in the MF-CA3 pathway. Dentate GC-specific deletion of *Igsf8* did not alter MF bouton number but reduced number and length of synaptic junctions at MF-CA3 synapses. Consistent with these morphological alterations, sEPSC frequency and amplitude in CA3 pyramidal neurons were reduced. These findings suggest that IgSF8 promotes MF-CA3 synapse development. In line with these observations, an immunohistochemical study demonstrated transient IgSF8 immunoreactivity in olfactory sensory neuron axon terminals during synaptogenesis (Ray and Treloar, 2012). Lesion-induced re-formation of synapses caused IgSF8 to reappear in terminals, whereas blocking olfactory activity prevented the disappearance of IgSF8 from mature synapses (Ray and Treloar, 2012). Similarly, the EWI-subfamily member IgSF3 transiently localizes to developing cerebellar GC axon terminals and inhibits axon growth, which might play a role in the timing of synapse formation (Usardi et al., 2017).

CSP interactome screening and affinity chromatography with the IgSF8 ectodomain identified the ECM protein TenR as a novel IgSF8 ligand at MF-CA3 synapses. The precise subcellular localization of TenR remains unclear. An electron microscopy study detected TenR in the extracellular space surrounding MF boutons but not in the synaptic cleft (Schuster et al., 2001), whereas proximity biotinylation identified TenR in the excitatory synaptic cleft of cultured neurons (Cijsouw et al., 2018). Perisomatic inhibitory synapses in the hippocampus of *Tnr* KO mice show decreased AZ length and number (Nikonenko et al., 2003), reminiscent of the morphological defects at *Igsf8* cKO MF-CA3 synapses, suggesting that the IgSF8-TenR interaction might stabilize synapses. Supporting this notion, synapse density is reduced in neuronal cultures lacking TenR (Geissler et al., 2013). Conversely, upregulated TenR protein levels following lesion-induced sprouting have been linked to axon targeting and synapse formation (Brenneke et al., 2004). Interactions of secreted ECM proteins and neuronal receptors play important roles in the development of synaptic connectivity (Yuzaki, 2018) and in the modulation of synaptic function and plasticity (Dityatev et al., 2010), but are poorly understood. IgSF8 also interacts with tetraspanins (Charrin et al., 2003; Stipp et al., 2001), which can promote synapse formation and function (Kopczynski et al., 1996) but remain poorly characterized at mammalian synapses (Murru et al., 2018).

In addition to defects in the main MF bouton, conditional deletion of IgSF8 strongly decreased the number of MF filopodia that target interneuron dendrites in SL (Acsády et al., 1998). IgSF8 localizes to neurites and growth cones in cultured neurons (Ray and Treloar, 2012). The cytoplasmic tail of IgSF8 binds Ezrin-Radixin-Moesin (ERM) proteins (Sala-Valdés et al., 2006) and alpha-actinin (Gordón-Alonso et al., 2012), both of which are linked to filopodia formation. TenR treatment induces the formation of actin-rich protrusions in cultured chick tectal neurons (Zacharias et al., 2002). Together, these observations suggest that an IgSF8-TenR interaction may similarly promote formation or stabilization of MF filopodia. Interestingly, TenR immunoreactivity is highly concentrated around parvalbumin-positive SL interneurons (Brenneke et al., 2004; Saghatelyan et al., 2001), which are among several interneuron targets of MF filopodia (Acsády et al., 1998; Szabadics and Soltesz, 2009). Filopodial synapse number and strength varies depending on SL interneuron type (Szabadics and Soltesz, 2009), suggesting that target cell identity plays a role in connectivity and composition of filopodial synapses. A small number of adhesion molecules has been implicated in regulation of filopodia number: Kirrel3 (Martin et al., 2015), CDH9 (Williams et al., 2011), and SynCAM1 (Park et al., 2016). Together with IgSF8, these and possibly other CSPs may act in combinations to regulate filopodial synapse connectivity and function.

MF filopodia-mediated feedforward inhibition controls CA3 excitability and output (Torborg et al., 2010). MF filopodia are highly plastic structures (Galimberti et al., 2006). Feedforward inhibition in CA3 microcircuits increases upon learning through enhanced connectivity between MF filopodia and interneurons, important for memory precision (Guo et al., 2018; Ruediger et al., 2011). Conversely, feedforward inhibition in CA3 microcircuits decreases with aging, resulting in CA3 hyperactivity (Villanueva-Castillo et al., 2017). Impaired MF filopodia-mediated feedforward inhibition may also contribute to circuit dysfunction in neurodevelopmental disorders (Martin et al., 2015), underscoring the importance of precise molecular control of CA3 microcircuit connectivity for cognitive function.

Dissecting the molecular composition of specific connections remains a major challenge. Sorting of identified projections (Poulopoulos et al., 2019) and synapses (Pfeffer et al., 2020; this study), imaging-based approaches (Micheva et al., 2010; Zhu et al., 2018), affinity purification (Selimi et al., 2009) and proximity biotinylation (Li et al., 2020; Uezu et al., 2016) approaches each have their advantages and disadvantages (Schreiner et al., 2017). A combination of approaches will be required to understand the molecular mechanisms that regulate the precise patterns of connectivity that are required for cognitive function.

## Acknowledgments

We thank Franck Polleux, Dietmar Schmucker, Pierre Vanderhaeghen, Anirvan Ghosh, Dan Dascenco, Luís Ribeiro, and Sara Calafate for critical reading of the manuscript and De Wit lab members for helpful discussion and comments. We thank Pier Andrée Penttila and Christèle Nkama (VIB-KU Leuven FACS Core), the VIB-KU Leuven BioImaging Core, Joris Vandenbempt and Brenda Luong for experimental help; Nicola Fattorelli for data visualization advice; Etienne Herzog and Matthew Holt for experimental advice. Leica SP8x confocal microscope was provided by InfraMouse (KU Leuven-VIB) through a Hercules type 3 project (ZW09-03). N.A. is supported by the Fundação para a Ciência e a Tecnologia (FCT, grant number SFRH/BD/128869/2017). D.C. is supported by NSF IOS Grant # 1755189, RWJ Foundation grant #74260, and Research for Life (WMRF) 2019/301. J.N.S. is supported by R01AG061787. J.d.W is supported by ERC Starting Grant (#311083), FWO Odysseus Grant, FWO Project grant G0C4518N, FWO EOS grant G0H2818N, and Methusalem grant of KU Leuven/Flemish Government.

## Author contributions

N.A., J.N.S., and J.d.W. conceived the study and designed experiments. N.A., S.N.S, J.V., G.C., V.R., J.t.B., L.T., S.P., K.M.V., N.V.G., D.C., and K.D.W. performed experiments and analyzed data. N.A. and J.d.W. wrote the paper with input from all authors. All authors contributed to and approved the final version.

## Declaration of interests

The authors declare no competing interests.

## Data availability

MS raw files and search results can be accessed via Proteome Exchange at PXD013492.

## METHODS

### CONTACT FOR REAGENT AND RESOURCE SHARING

Further information and requests for resources and reagents should be directed to and will be fulfilled by the Lead Contact, Joris de Wit

### EXPERIMENTAL MODEL AND SUBJECT DETAILS

#### Animals

All animal experiments were conducted according to the KU Leuven ethical guidelines and approved by the KU Leuven Ethical Committee for Animal Experimentation (approved protocol numbers ECD P037/2016, P014/2017 and P062/2017). Mice were maintained in a specific pathogen-free facility under standard housing conditions with continuous access to food and water. Mice used in the study were 1-8 weeks old and were maintained on a diurnal 12-hour light/dark cycle. Wild-type (WT) C57BL/6J mice were obtained from JAX. The *Igsf8* conditional knock out (cKO) mouse line was obtained from the Riken Institute (Inoue et al., 2012) while the *Rpb4-Cre* line was obtained from GENSAT. Genotypes were regularly checked by PCR analysis. For euthanasia, animals were either anesthetized with isoflurane and decapitated or injected with an irreversible dose of ketamine-xylazine.

#### Neuronal Cultures

Hippocampal neurons were cultured from E18 *Igsf8* conditional knock out mice and plated on poly-D-lysine (Millipore) and laminin (Invitrogen)-coated glass coverslips (Glaswarenfabrik Karl Hecht). Neurons were maintained in Neurobasal medium (Invitrogen) supplemented with B27, glucose, glutamax, penicillin/streptomycin (Invitrogen), and 25 µm β-mercaptoethanol (Sigma-Aldrich). To knock out *Igsf8*, neurons were infected with low-titer lentivirus expressing mCherry or mCherry-2A-Cre two days after plating and collected at day *in vitro* 10.

#### Cell Lines

HEK293T-17 human embryonic kidney cells (available source material information: fetus) were obtained from American Type Culture Collection (ATCC) cat# CRL-11268. HEK293T-17 cells were grown in DMEM (Invitrogen) supplemented with 10% FBS (Invitrogen) and penicillin/streptomycin (Invitrogen).

### METHODS DETAILS

#### Plasmids

For the ELISA-based interactome assay we used recombinant AP- or Fc-tagged constructs containing human or mouse extracellular domain (ectodomain) protein sequences derived from the appropriate entry in the UNIPROT data base (Uniprot.org). Cloning boundaries and cloning strategy can be found elsewhere (Ozgul et al., 2019; Ranaivoson et al., 2019). Briefly, DNA sequences encoding the ectodomains of transmembrane and GPI-anchored CSPs, or the entire protein sequence in the case of secreted proteins, lacking the signal peptide, were cloned in frame using the 5’ NotI and 3’ XbaI restriction sites of a modified pCMV6-XL4 expression vector. Fc fusion proteins contain a leader peptide (PLP, prolactin leader peptide) followed by a N-terminal FLAG tag, ectodomain of interest, a 3CPro cleavage site and the dimeric human Fc domain. Similarly, AP fusion proteins contain a leader peptide, a FLAG tag and ectodomain of interest. C-terminal to the ectodomain there is a human AP followed by a His_6_ tag. For cloning Neurexin ectodomains we used DNA sequences of rat Neurexin1α-(-S4), mouse Neurexin1β-(- S4), mouse Neurexin3α-(-S4) (cDNA gift from Ann Marie Craig’s lab) and rat Neurexin3β-(-S4) (cDNA from Addgene plasmid #58269 from Peter Scheiffele’s lab). All constructs were verified by DNA sequencing. The fact that AP is enzymatically active provides a simple way to test expression and measure accurate protein concentration. The 146 recombinant AP- or Fc-tagged constructs used for this ELISA interaction screen were secreted into the media after transient transfection, and used in the assay without further purification. Four CSPs were not included in the ELISA-based interactome assay: CNTN1AP is not expressed in the absence of CNTN1; LRP1 and VCAN are very large proteins and costly to synthesize; and RPTPε has a very small ectodomain making potential interactions difficult to detect using this assay.

Full-length cDNAs encoding mouse IgSF8 (BC048387) and Tenascin-R (BC138043) were purchased from Origene and Source Bioscience, respectively. Both cDNAs were cloned into the pEGFP-N1 vector (Clontech) to place a C-terminal EGFP tag. Full-length cDNA encoding mouse Brevican (BC052032), lacking the signal peptide, was purchased from Source Bioscience and cloned into the Fc-tagged construct.

To generate pAAV-hsyn1-mGFP-T2A-Cre we amplified EGFP-T2A-Cre by PCR from pRetroX-GFP-T2A-Cre (Addgene #63704) and inserted this fragment between restriction sites BamHI and HindIII of pAAV-hSyn1-ChR2-EGFP (Addgene #58881). After, mGFP was PCR-amplified from FUGW-mGFP-T2A-HA-GPR158 (not published) and replaced by EGFP in pAAV-hsyn1-EGFP-T2A-Cre. As a control vector, pAAV-hsyn1-mGFP was generated by inserting PCR-amplified mGFP between restriction sites BamHI and HindIII of pAAV-hSyn1-ChR2-EGFP (Addgene #58881). The plasmid pAAV-Ef1a-DIO-hChR2(E123T/T159C)-EYFP was obtained from Addgene (#35509).

#### Isolation of MF-CA3 synaptosomes

For initial characterization and comparison of crude MF-CA3 synaptosomes and standard hippocampal synaptosomes, we followed the protocol as described in Taupin et al. (Taupin et al., 1994). Briefly, hippocampi were dissected quickly in ice-cold Hank’s Balanced Salt Solution (HBSS) and homogenized in homogenization buffer (0.32 M sucrose, 5 mM Trizma Base, 1 mM MgCl2, pH 7.4) with protease inhibitors (pepstatin A, leupeptin, aprotinin and PMSF) using a Dounce homogenizer. Homogenate was spun at 1,000 x g to pellet MF-CA3 synaptosomes with nuclei and large cell debris in pellet 1 (P1), while general small synaptosomes are in the supernatant (S1). P1 and S1 fractions were loaded on Percoll gradients (Thermo-Fisher) and spun at 32,500 x g at 4°C for 7 minutes to separate MF-CA3 and standard synaptosomes from myelin, nuclei or mitochondria, resulting in PI (crude MF-CA3) and SI (small hippocampal) synaptosome fractions. To prepare MF-CA3 synaptosomes for sorting, we adapted the protocol of *Taupin et al.* using 10-12 post-natal day 28 (P28) WT mice per experiment. Hippocampi were dissected quickly in ice-cold HBSS and homogenized in homogenization buffer (0.32 M sucrose, 4 mM Hepes, 1 mM MgCl2, pH 7.4) with protease inhibitors (pepstatin A, leupeptin, aprotinin and PMSF) using a Dounce homogenizer. After, the homogenate was filtered through a series of cell strainers (100µm, 70 µm and 30 µm) and spun at 1,000 x g for 10 minutes at 4°C to prepare P1, which includes large MF-CA3 synaptosomes, and S1. P1 was washed once by re-suspension in homogenization buffer and re-centrifuged as above. Supernatants S1 were pooled and spun at 15,000 x g for 20 minutes at 4°C to prepare post-nuclear pellet (P2) synaptosomes. P1 and P2 were re-suspended in PBS and myelin was depleted from both fractions according to the instructions of a commercial myelin-removal kit (Miltenyi Biotec), before sorting MF-CA3 synaptosomes from myelin-depleted P1.

#### Fluorescence-activated synaptosome sorting

Myelin-depleted MF-CA3 synaptosomes were prepared as described above. After, MF-CA3 synaptosomes were labeled in non-permeabilizing conditions in PBS with an anti-Nectin 3 monoclonal antibody (1:50; Hycult Biotech) directly conjugated with CF488A fluorophore (Sigma-Aldrich) after depletion of bovine serum albumin present in the antibody storage solution, using an antibody clean-up kit (Thermo-Fisher). Immediately before running the samples in the cytometer, FM4-64 (Thermo-Fisher) was added to myelin-depleted MF-CA3 synaptosomes in order to label all membrane-containing material. FM4-64/Nectin 3-488-double positive MF-CA3 synaptosomes were sorted on a BD FACS Aria III using an 85μm nozzle at 45psi. The pressure differential was set to 2 to minimize sheer stress. Nectin 3-488 was excited using a 488nm 13mW laser and detected with a 530/30 band pass filter. FM4-64 was excited using a 561nm 30mW laser and detected with a 720/40 band pass filter. Unstained and single stained CF488A and FM4-64 synaptosomes were analyzed to calculate autofluorescence and signal above background, respectively, and Nectin 3-488 and FM4-64 gates were set accordingly. Synaptosomes were identified by back-gating on fluorescence using FSC and SSC, with SSC set on a 5 decade log scale. FM4-64/Nectin 3-488-double positive MF-CA3 synaptosomes were analyzed on the BD FACS Aria III post-sort to quantify the purity of the sorted population.

#### Protein expression

For the ELISA-based assay, proteins were expressed in Expi293 suspension adapted cells (ThermoFisher Scientific). These cells were cultured in Expi293™ Expression Medium and transfection was achieved by mixing 2µg of cDNA with 6µg of polyethylenimine (PEI) and adding the mixture to two millions cells in 0.8mL of medium. Conditioned media was harvested 4-6 days after transfection and used within two days. Fc constructs were characterized by SDS-PAGE to ensure protein integrity and confirm the MW of the sample. Fc fusion proteins were quantified by Western blot using a known quantity of Fc-only protein. AP fusion proteins are quantified using calf intestinal alkaline phosphatase (CIP, New England Biolabs) activity. Briefly, the activity of 20 µL of conditioned medium is compared with the activity of 1 µL of CIP at room temperature after 5 minutes in 50 µL of reaction in Expi293 medium. All of the AP-fusion proteins expressed at levels >1 U/µL of CIP (where 1 U=10 pg of purified CIP).

#### ELISA protocol

An ELISA-based assay was used to test the binding between ectodomain-Fc and ectodomain-AP fusions. Each data point was reproduced in three independent experiments, and each ectodomain was tested in both orientations. Fifteen μl of a solution at 3 µg/mL of mouse anti-AP (IgGAb-1 clone 8B6.18 Thermo Fisher Scientific; Waltham, MA) in 1X PBS was added to each well of 384-well plates using an automated multichannel pipette (Viaflo Assist, Integra), sealed and incubated overnight at 4°C. The following day, plates were washed, and 5% dry milk was added as a blocking agent, which was removed after 1 hr at room temperature using an automated microplate washer (HydroSpeed, Tecan). Subsequently, to each well, 9 μl of ecto-AP conditioned medium containing 2 μL of monoclonal mouse anti-human IgG1-HRP (2 μg/ml; Serotec; Raleigh, NC) was added using an automated plate copier (Viaflo96, Integra) along with 7 μl of ecto-Fc culture medium. Plates were sealed and incubated for 4 hr at room temperature in the dark. Plates were subsequently washed, and 15 µL 1-Step Ultra TMB-ELISA HRP substrate was added using an automated multichannel pipette (Viaflo Assist, Integra); after 1 hr incubation at room temperature, the absorbance at 650 nm was recorded with a Perkin Elmer EnSpire 2300 multimode plate reader. Finally, plates were scanned to obtain matching images of the 650 nm reading. Known interactors such as NRXNs/NLGNs and IgLONs/IgLONs, were among the tested pairs and worked well as positive controls. Negative control bait (AP only protein) was used to detect false positive interactions. Background values necessary to quantify the positive reactions were obtained by averaging at least 30 contiguous “blank” wells (within a typical plate, the mean background Abs650 for 100 negative wells was 0.07 ± 0.027). Dynamic range of a typical plate was ∼50FOB.

#### Cell Surface Binding Assay

HEK293T cells were grown in Dulbecco’s modified Eagle’s medium (DMEM) (Invitrogen) supplemented with 10% FBS (Invitrogen), penicillin/streptomycin and transfected with EGFP, myc-LRRTM2, IgSF8-EGFP and Tenascin-R-EGFP plasmids using Fugene6 (Promega). Twenty-four hours after transfection, the cells were incubated with Fc, Neurexin1β-Fc, IgSF8-Fc or Tenascin-R-Fc proteins (10 µg/ml) in DMEM supplemented with 20 mM Hepes (pH 7.4) for 1 hour at room temperature (RT). Following two brief washes in DMEM/20 mM Hepes pH 7.4, cells were fixed and immunostained using mouse anti-GFP (Santa Cruz), mouse anti-c-myc (Santa Cruz) and the Cy3-conjugated anti-Fc antibody (Jackson ImmunoResearch). Fluorophore-conjugated secondary antibodies were from Jackson ImmunoResearch or Invitrogen. The cells were imaged with a Leica SP8 confocal microscope (Leica Microsystems) using a 63X objective.

#### Fc-Protein Purification

IgSF8-Fc and Tenascin-R-Fc proteins were produced by transient transfection of HEK293T cells using PEI (Polysciences). Six hours after transfection, media was changed to OptiMEM (Invitrogen) and harvested 5 days later. Conditioned media was centrifuged, sterile-filtered and run over a fast-flow Protein-G agarose (Thermo-Fisher) column. After extensive washing with wash buffer (50 mM Hepes pH 7.4, 300 mM NaCl and protease inhibitors), the column was eluted with Pierce elution buffer. Eluted fractions containing proteins were pooled and dialyzed with PBS using a Slide-A-Lyzer (Pierce) and concentrated using Amicon Ultra centrifugal units (Millipore). The integrity and purity of the purified ecto-Fc proteins was confirmed with SDS-PAGE and Coomassie staining, and concentration was determined using Bradford protein assay.

#### Affinity Chromatography

Affinity chromatography experiments were performed as previously described (45). P1 crude MF-CA3 synaptosomes were prepared as described above from mouse brains, without the myelin removal step. For the preparation of P2 crude synaptosome extracts, ten P21-22 rat brains were homogenized in homogenization buffer (4 mM Hepes pH 7.4, 0.32 M sucrose and protease inhibitors) using a Dounce homogenizer. Homogenate was spun at 1,000 x g for 10 minutes at 4°C. Supernatant was spun at 14,000 x g for 20 minutes at 4°C. P1 containing crude MF-CA3 synaptosomes and P2 crude synaptosomes were re-suspended in Extraction Buffer (50 mm Hepes pH 7.4, 0.1 M NaCl, 2 mM CaCl2, 2.5 mM MgCl2 and protease inhibitors), extracted with 1% Triton X-100 for 2 hours and centrifuged at 100,000 x g for 1 hour at 4°C to pellet insoluble material. Fast-flow Protein-A sepharose beads (GE Healthcare) (250 µl slurry) pre-bound in Extraction Buffer to 100 µg human Fc or IgSF8-ecto-Fc were added to the supernatant and rotated overnight at 4°C.

Beads were packed into Poly-prep chromatography columns (BioRad) and washed with 50 ml of high-salt wash buffer (50 mM HEPES pH 7.4, 300 mM NaCl, 0.1 mM CaCl2, 5% glycerol and protease inhibitors), followed by a wash with 10 ml low-salt wash buffer (50 mM HEPES pH 7.4, 150 mM NaCl, 0.1 mM CaCl2, 5% glycerol and protease inhibitors). Bound proteins were eluted from the beads by incubation with Pierce elution buffer and TCA-precipitated overnight. The precipitate was re-suspended in 8 M Urea with ProteaseMax (Promega) per the manufacturer’s instruction. The samples were subsequently reduced by 20-minute incubation with 5mM TCEP0 (tris(2carboxyethyl)phosphine) at RT and alkylated in the dark by treatment with 10 mM Iodoacetamide for 20 additional minutes. The proteins were digested overnight at 37°C with Sequencing Grade Modified Trypsin (Promega) and the reaction was stopped by acidification. Mass spectrometry analysis was performed as before (Savas et al., 2014).

#### Mass spectrometry

Sorted MF-CA3 synaptosomes and P2 synaptosomes were collected and protein extracted, precipitated and loaded in SDS-PAGE gels. Gels were further stained with InstantBlue Protein Stain (expedeon) according to instructions. Protein lanes were cut in four sections using clean material under the laminar flow. Each section was further cut in approximately 1mm3 cubes, transferred to individual microcentrifuge tubes and incubated three times in 30% ethanol at 60°C for 20 minutes to wash excess InstantBlue Protein Stain. To prepare samples for mass spectrometry, each gel section was brought to RT and submerged in 100 µl buffer containing ammonium bicarbonate at 50 mM and TCEP at 10 mM for cystine reduction. The gel samples were incubated in this solution at 37°C for one hour whilst vortexing. The liquid was removed and replaced with 100 µl of a new solution of 50 mM ammonium bicarbonate and 50 mM iodoacetamide for cystine alkylation. The gel samples were then incubated in the solution at RT in the dark for 45 minutes. Then, the liquid was removed and replaced with 100 µl of 50 mM ammonium bicarbonate and 50 mM TCEP to quench alkylation reaction at RT for 30 minutes. The liquid was then removed, and the gel sections were washed twice with a solution of 50 mM ammonium bicarbonate. The gel sections were then each submerged by a 100 µl solution of 50 mM ammonium bicarbonate and 1 µg of Trypsin and incubated overnight at 37°C with vortexing for protein digestion to peptides. The next day, the liquid from the digested gel samples was collected in new microcentrifuge tubes. Then the gel slices were treated a total of three times with a solution of 50% acetonitrile, 5% formic acid for 20 minutes followed by collection of the liquid in the new tube. The tubes now containing the extracted peptides from the gel slices were dried by vacuum centrifugation. The dried peptides were then re-suspended in 200 µl of a solution of 0.5% trifluoroacetic acid and desalted on C18 resin spin columns (Pierce) using 5% acetonitrile, 0.5% trifluoroacetic acid to wash and 80% acetonitrile, 0.5% formic acid buffer to elute. These desalted samples were then dried by vacuum centrifugation and resuspended in LC-MS/MS sample buffer of 5% acetonitrile, 1% formic acid.

For mass spectrometry data acquisition from the samples, 3 µg of peptides corresponding to each of the four gel sections of each of the two samples from each experiment was injected into a Thermo Orbitrap Fusion Mass Spectrometer equipped with an Ultimate 3000 RSLCnano liquid chromatography. Peptides were loaded onto a vented Acclaim Pepmap 100, 75 µm x 2 cm, nanoViper trap column and subsequently separated using a nanoViper analytical column (Thermo-164570, 3µm, 100Å, C18, 0.075mm, 500mm) and electrosprayed with stainless steel emitter tip assembled on the Nanospray Flex Ion Source with a spray voltage of 2000V. Buffer A contained H2O with 5% ACN and 0.125% FA, and buffer B contained H2O with 95% ACN and 0.125% FA. The chromatographic run was for 160 minutes in total with the following profile of percent buffer B: 0 to 8% for 6 minutes, to 24% for 64 minutes, to 36% for 20 minutes, to 55% for 10 minutes, to 95% for 10 minutes, held at 95% for 20 minutes, then brought down to 2% over 1 minute, and held at 2% for 29 minutes. Additional mass spectrometry parameters include: Ion transfer tube temp at 300°C, positive spray voltage at 2500V, and top speed with a cycle time of 3 seconds. MS1 scan in Orbitrap with 120K resolution, scan range = 300-1800 (m/z), max injection time = 50ms, AGC target = 100,000, microscans = 1, RF lens = 60%, and datatype = centroid. MIPS mode = peptide, included charge states = 2-8 (reject unassigned). Dynamic exclusion enabled with n = 1 for 30 second exclusion duration at 5 ppm for high and low. Precursor selection decision for MS2 scan in the ion trap set at most intense with a 1000 minimum intensity threshold, isolation window = 1.6, scan range = auto normal, first mass = 100, collision energy 32% HCD, IT scan rate = rapid, max injection time at 50ms, AGC target = 10,000, datatype = centroid, inject ions for all available parallelizable time.

Spectrum raw files from samples were extracted into ms1 and ms2 files using in-house program RawConverter (http://fields.scripps.edu/downloads.php) (He et al., 2015), and the tandem mass spectra were searched against UniProt mouse protein database (downloaded on 03-25-2014) (The UniProt Consortium 2015) and matched to sequences using the ProLuCID/SEQUEST algorithm (ProLuCID ver. 3.1) (Eng et al., 1994; Xu et al., 2015) with 50 ppm peptide mass tolerance for precursor ions and 600 ppm for fragment ions. The search space included all fully and half-tryptic peptide candidates that fell within the mass tolerance window with no miscleavage constraint, assembled and filtered with DTASelect2 (ver. 2.1.3) (Cociorva et al., 2007; Tabb et al., 2002) through Integrated Proteomics Pipeline (IP2 v.3, Integrated Proteomics Applications, Inc., CA, USA http://www.integratedproteomics.com). To estimate peptide probabilities and false-discovery rates (FDR) accurately, we used a target/decoy database containing the reversed sequences of all the proteins appended to the target database. Each protein identified was required to have a minimum of one peptide of minimal length of five amino acid residues and within 10 ppm of the expected m/z. However, this peptide had to be an excellent match with an FDR less than 0.001 and at least one excellent peptide match. After the peptide/spectrum matches (PSM) were filtered, we estimated that the protein FDRs were ≤ 1% for each sample analysis.

To identify the unique protein composition of MF-CA3 synapses, first we filtered our protein list on the criteria that a candidate protein had to have at least three peptide identifications among the six samples. Then Log2 fold change of sorted MF-CA3 synaptosomes versus the P2 synaptosome background was calculated on the basis of the measured normalized spectral abundance factor (NSAF). This measure normalizes spectral counts for a protein based on the total number of spectra in the run, the length of the protein, and the total number of peptides that comprise it (Florens et al., 2006; Zybailov et al., 2006). Proteins that were not identified in a sample after filtering had their NSAF values imputed as 0 for all calculations. Proteins were determined significant if the Student’s T-Test p value was <0.05 among the three replicates. A q value was calculated based on Benjamini-Hochberg procedure and the highest ranked protein with a q value <0.05 was used as a cutoff for determining high-confidence measurements at a 5% FDR.

#### Gene ontology analysis

To find cellular component terms overrepresented in sorted MF-CA3 synaptosomes or P2 synaptosomes we used the statistical overrepresentation test in the Panther Classification System using the mouse genome as reference (http://www.pantherdb.org/) (Mi et al., 2019). To analyze the distribution of the MF-CA3 synaptic proteome within major cellular components we used the functional classification tool in Panther. To select CSPs among the MF-CA3 synaptic proteome we manually queried UniProt (https://www.uniprot.org/) for annotated proteins with transmembrane and extracellular domains, as well as secreted and extracellular matrix proteins. CSP functional class and protein domains were also manually examined using UniProt and known literature in PubMed (https://www.ncbi.nlm.nih.gov/pubmed/).

#### Fc Pulldown Assays

For pulldown assays on HEK293T cells, cells were grown in 10 cm dishes in DMEM (Invitrogen) supplemented with 10% FBS (Invitrogen) and penicillin/streptomycin, and transfected with EGFP, IgSF8-EGFP or Tenascin-R-EGFP expression constructs using Fugene6 (Promega). Twenty-four hours after transfection, the media was changed to OptiMEM (Invitrogen) for 2 hours. Cells were then lysed in 1 ml ice-cold RIPA buffer (20 mM Tris-HCl pH 7.5, 150 mM NaCl, 5 mM EDTA, 1% Triton X-100 and protease inhibitors (Roche)) for 1 hour at 4°C on a rocking platform. Lysates were spun at 13,000 rpm for 30 minutes at 4°C. Three µg of human Fc, IgSF8-Fc or Tenascin-R-Fc was added to 1 ml of supernatant and rotated overnight at 4°C. Protein-A agarose beads (50 µl slurry) were added and rotated for 1 hour at 4°C. Beads were washed 3 times in cold RIPA buffer and once in PBS, boiled in 50 µl 2X sample buffer and analysed by Western blotting using a mouse anti-GFP primary antibody (Santa Cruz) and HRP-conjugated goat anti-mouse IgG secondary antibody (Thermo-Fisher).

#### Subcellular Fractionation

Synaptic fractionation was based on a previously described method (Carlin et al., 1980). In brief, ten P21 rat brains were homogenized in 12 ml per brain with homogenization buffer (0.32 M sucrose, 4 mM Hepes pH 7.4, 1 mM MgCl2 and protease inhibitors) (homogenate), centrifuged at 1,500 x g for 15 minutes, and the supernatant was collected (post nuclear supernatant). The supernatant was then centrifuged at 18,000 x g for 20 minutes, and the resulting supernatant (cytosol) and pellet (crude membrane) collected. The pellet was re-suspended in homogenization buffer and loaded onto 0.85 M/1.0 M/1.2 M discontinuous sucrose gradients and centrifuged at 78,000 x g for 120 minutes. The material at the 1.0 M/ 1.2 M interface was collected (synaptosome). Triton X-100 was added to 0.5% and extracted at 4°C by end-over-end agitation for 20 minutes. The extract was centrifuged at 32,000 x g for 20 minutes, the supernatant collected (soluble synaptosome/Triton-soluble fraction) and the pellet was re-suspended in Homogenization Buffer, loaded onto a 1.0 M/1.5 M/2.0 M sucrose gradient, and centrifuged at 170,000 x g for 2 hours. Material was collected at the 1.5 M/2.0 M interface (PSD). 0.5% Triton X-100 was added and detergent-soluble material extracted at 4°C by end-over-end agitation for 10 minutes. Lastly, the extract was centrifuged at 100,000 x g for 20 minutes and the pellet re-suspended in homogenization buffer (purified PSD/Triton-insoluble fraction). Primary antibodies used were the following: mouse anti-PSD95 (Thermo-Fisher), rabbit ant-Synaptophysin (Sigma-Aldrich) and goat anti-IgSF8 (R&D Systems). HRP-conjugated secondary antibodies were from Thermo-Fisher.

#### Immunocytochemistry

Crude MF-CA3 (PI fraction) and small hippocampal (SI fraction) synaptosomes were spun at 15,000 x g for 20 min at 4°C after being collected, re-suspended in PBS, transferred to PDL-coated 8-well Nunc Lab-Tek chamberslides (Thermo-Fisher) and incubated at 4°C on a shaker for 90 minutes to allow synaptosomes to settle. After, synaptosomes were fixed in 2% paraformaldehyde and stained. Briefly, synaptosomes were blocked and permeabilized in 3% BSA, 0.1% Saponin in PBS (BSA-BLOCK) at RT for 30 minutes and incubated with primary antibodies in BSA-BLOCK at 4°C overnight. After, synaptosomes were washed three times with PBS, incubated with secondary antibodies at RT for 1 hour, washed three times with PBS, incubated with Hoechst dye at RT for 5 minutes, washed again and mounted with Mowiol-4-88 (MilliporeSigma Calbiochem). For live labeling, crude MF-CA3 synaptosomes were incubated in non-permeabilizing conditions with primary antibodies diluted in 3% BSA in PBS before blocking, permeabilization and staining of intracellular epitopes. In sorting experiments, a sample of myelin-depleted pre-sorting P1 MF-CA3 synaptosomes was collected before sorting. Remaining sample was sorted and spun at 1,000 x g for 10 minutes at 4°C to pellet sorted MF-CA3 synaptosomes. Finally, sorted MF-CA3 synaptosomes were re-suspended in PBS and stained in permeabilizing conditions as above, in parallel to myelin-depleted pre-sorting MF-CA3 synaptosomes. Primary antibodies were the following: rat anti-Nectin3 (Hycult Biotech), mouse anti-GluK5/KA2 (NeuroMab), rabbit anti-Synaptoporin (Synaptic Systems), guinea pig anti-VGluT1 (Millipore) and mouse anti-PSD95 (Thermo-Fisher). Fluorophore-conjugated secondary antibodies were from Jackson ImmunoResearch or Invitrogen. Confocal images were acquired using a Leica SP8 confocal microscope (Leica Microsystems) and analyzed with Fiji (Schindelin et al., 2012).

#### Immunohistochemistry

P28 WT mice were anesthetized by intraperitoneal injection with a lethal dose of 1 μl/g Xylazine (VMB Xyl-M 2%), 2μl/g Ketamine (Eurovet Nimatek, 100 mg/ml) and 3 μl/g 0.9% saline. Next, mice were transcardially perfused with 4% paraformaldehyde, in PBS. Brains were dissected, postfixed in 4% paraformaldehyde in PBS at 4°C for one hour and cryopreserved in 30% sucrose overnight at 4°C. After, perfused brains were embedded in Tissue-Tek O.C.T. compound (Sakura) and frozen in isopentane at −55 to −65°C. For fresh frozen immunohistochemistry, mice were euthanized using isoflurane (Halocarbon). Next, brains were quickly dissected, embedded in Tissue-Tek O.C.T. compound (Sakura), and frozen in isopentane at −55 to −65°C. To prepare hippocampal sections we used the cryostats (NX70, Thermo Fisher) and CM3050 S (Leica Biosystems). Frozen brains were cut and 16-20 μm thick coronal sections were collected on SuperFrost Ultra Plus adhesion slides (Thermo-Fisher). The fresh frozen sections were postfixed with 1:1 acetone-methanol for 5 minutes at −20°C and quickly washed in PBS before staining.

In some cases, heat-induced antigen retrieval (H-AR) was done before immunostainings in sections of paraformaldehyde perfused brains. For H-AR, sections were submerged in sodium citrate buffer containing 0.05% Tween20 and 10 mM Sodium Citrate pH of 6.0 and heated in either a microwave up to boiling, for 30 seconds, twice for 15 seconds and once for 10 seconds with a 1-2 minutes cooling period at RT between heating; or in the 2100 Antigen Retriever pressure cooker (Aptum Biologics Ltd) for 25 minutes up to 120°C. After H-AR, sections were washed in PBS. Both fresh frozen and perfused sections were then permeabilized at RT for 20 minutes in PBS containing 0.05% Triton X-100 (Sigma). Sections were then blocked at RT for 2 hours in PBS containing 10% normal horse serum (NHS), 0.5% Triton X-100, 10% glycine 2 M, 0.02% gelatin and, in the case of using mouse primary antibodies, 1:50 donkey anti-mouse IgG antigen-binding fragments (Jackson ImmunoResearch). Sections were then washed in PBS-0.5%Triton X-100 at RT and incubated at 4°C overnight with primary antibodies in PBS containing 5% NHS, 0.5% Triton X-100 and 0.02% gelatin. Afterwards, sections were washed in PBS-0.5%Triton X-100 at RT before a 2 hour incubation at RT with secondary antibodies in PBS containing 5% NHS, 0.5% Triton X-100 and 0.02% gelatin. Before mounting coverslips with Mowiol-4-88 (MilliporeSigma Calbiochem), sections were washed in PBS-0.5% Triton X-100. Primary antibodies were the following: rat anti-Nectin3 (Hycult Biotech), mouse anti-GluK5/KA2 (NeuroMab), rabbit anti-Synaptoporin (Synaptic Systems), guinea pig anti-VGluT1 (Millipore), rabbit anti-Synapsin3 (Synaptic Systems), rabbit anti-Afadin (Thermo-Fisher), chicken anti-ZnT3 (Synaptic Systems), rabbit anti-mGluR2/3 (Merck Millipore), rabbit anti-CRTAC1 (Merck Millipore), goat anti-IgSF8 (R&D Systems), mouse anti-PSA-NCAM (Merck Millipore), rat anti-NCAM2 (R&D Systems), sheep anti-ISLR2 (R&D Systems), goat anti-NEGR1/Kilon (R&D Systems), mouse anti-Neuronal pentraxin 1 (BD Biosciences), goat anti-NRP1 (R&D Systems), goat anti-NRP2 (R&D Systems), sheep anti-Noelin (R&D Systems), goat anti-Plexin-A3 (Thermo-Fisher), rabbit anti-ROBO2 (Aviva Systems), goat anti-Neogenin (R&D Systems), sheep anti-SALM2 (R&D Systems), rabbit anti-FAM171A2 (ThermoFisher), mouse anti-Tenascin-R (R&D Systems), sheep anti-Contactin 1 (R&D Systems), goat anti-CD200 (R&D Systems), goat anti-ICAM5 (Novus), goat anti-Kit/SCFR (R&D Systems), mouse anti-LSAMP (Developmental Studies Hybridoma Bank), mouse anti-Trk-C (R&D Systems), goat anti-Plexin-A1 (R&D Systems) and goat anti-PTPRS (R&D Systems). Fluorophore-conjugated secondary antibodies were from Jackson ImmunoResearch or Invitrogen.

For free-floating immunohistochemistry, mouse anesthesia, perfusion and brain dissection were performed as previously described. Postfixation was done overnight at 4°C in PBS containing 4% paraformaldehyde. After, brains were washed three times with PBS at RT and embedded in 3% agarose. After, 80 μm thick sections were collected in PBS containing 0.2% NaN3 using the Vibrating Microtome 7000 (Campden Instruments LTD). Vibratome sections were washed at RT in PBS-0.5% Triton X-100. Next, sections were blocked at 4°C overnight in PBS containing 10% NHS, 0.5% Triton X-100, 0.5 M glycine and 0.2% gelatin. Thereafter, sections were incubated overweekend with primary antibodies in PBS containing 5% NHS, 0.5% Triton X-100 and 0.2% gelatin. Sections were then washed with PBS-0.5%Triton X-100 and incubated with the secondary antibodies in PBS containing 5% NHS, 0.5% Triton X-100 and 0.2% gelatin, at 4°C overnight. Subsequently, sections were washed in PBS-0.5%Triton X-100. To stain nuclei sections were incubated in PBS containing Hoechst (1:200) for 10 minutes at RT. Finally, sections were washed in PBS, collected on a microscope slide and coverslips were mounted with Mowiol 4-88 (MilliporeSigma Calbiochem). Primary antibodies were the following: chicken anti-GFP (Aves Labs), rabbit anti-Synaptoporin (Synaptic Systems) and goat anti-IgSF8 (R&D Systems). Fluorophore-conjugated secondary antibodies were from Jackson ImmunoResearch or Invitrogen. Confocal images were acquired using a Leica SP8 confocal microscope (Leica Microsystems) and analyzed with Fiji (Schindelin et al., 2012).

#### Western blots

Sorted MF-CA3 synaptosomes were collected into sorting tubes containing 4x lysis buffer (40mM Hepes, 600mM NaCl, 4% NP-40, 4% sodium deoxycholate, 0.4% SDS and 20mM EDTA). P2 synaptosomes were re-suspended in 2x lysis buffer and both samples incubated on ice for 30 minutes and further rotated at 4°C for 1 hour. To precipitate proteins, 20% (v/v) of freshly prepared TCA was added to the lysate of both samples. After, lysates were vortexed and incubated on ice overnight in the cold room. Finally, precipitates were spun at 13,000 g at 4°C for 30 minutes, washed three times with ice-cold acetone and air-dried. Protein precipitates were then re-suspended in 4x Laemmli buffer (8% SDS, 40% glycerol, 20% β-mercaptoethanol, 0.01% bromophenol blue and 250mM Tris HCl pH 6.8), pH-adjusted with 1.5M Tris HCl pH 8.8, spun, boiled at 95°C for 5 minutes and loaded in a gel for SDS-PAGE. Primary antibodies were the following: mouse anti-PSD95 (Thermo-Fisher), rabbit ant-Synaptophysin (Sigma-Aldrich), rabbit anti-Synaptoporin (Synaptic Systems), rabbit anti-Nectin3 (Abcam), goat anti-MBP (Santa Cruz), rabbit anti-Histone H3 (Cell Signaling Technology), goat anti-IgSF8 (R&D Systems), mouse anti-TenR (R&D Systems), rabbit anti-BRINP2 (Atlas Antibodies), sheep anti-CNTN1 (R&D Systems), goat anti-ICAM5 (R&D Systems), sheep anti-ISLR2 (R&D Systems), goat anti-Kit/SCFR (R&D Systems), goat anti-NEGR1 (R&D Systems), goat anti-Neogenin (R&D Systems), goat anti-NRP1 (R&D Systems), goat anti-Plexin-A1 (R&D Systems), rabbit anti-PTPRD (Novus), goat anti-PTPRS (R&D Systems), rabbit anti-ROBO2 (Abcam) and sheep anti-Teneurin-4 (R&D Systems). HRP-conjugated secondary antibodies were from Thermo-Fisher.

#### Direct Binding Assay

For direct binding assays of IgSF8 to Tenascin-R and Brevican, 1 µg recombinant His-tagged mouse IgSF8 (Sino Biological) was incubated in 1 ml binding buffer (10 mM HEPES pH 7.4, 150 mM NaCl, 2 mM CaCl2, 1 mM MgCl2 and 0.1% Tween-20) with equimolar amounts of control Fc protein (Jackson ImmunoResearch), purified Tenascin-R-Fc or Brevican-Fc and rotated end-over-end for 1 hour at RT. Protein A/G agarose beads (100 µl slurry; Santa Cruz Biotechnology) were added and rotated end-over-end for 1 hour at 4°C. Beads were washed 4x in binding buffer and 1x in PBS and boiled in 50 µl 2x sample buffer. Samples were analyzed by Western blot.

#### Transmission Electron Microscopy

To analyze crude MF-CA3 synaptosomes, the P1 fraction collected from 10 mice was fixed with 2% glutaraldehyde, 4% paraformaldehyde, 0.2% picric acid in 0.1M PB pH 7.40, gelatin embedded, and 100 μm vibratome sections were incubated in 1% osmium tetroxide, 1.5% potassium hexacyanoferrate in 0.1M cacodylate buffer, and then dehydrated with ethanol. Subsequently, sections were contrasted in 2% uranyl acetate, and en block lead acetate, washed, and embedded in Epon. Samples were sectioned as 70 nm, collected on copper grids and examined with a Jeol Jem 1400 electron microscope.

To analyze morphological changes in *Rbp4-Cre:Igsf8* cKO double transgenic mice, P30 littermates (*Rbp4-Cre^+^:Igsf8^+/+^ and Rbp4-Cre^+^:Igsf8^f/f^*) were transcardially perfused with 4% paraformaldehyde, 2.5% glutaraldehyde, 0.2% picric acid in 0.1M PB. Brains were removed, embedded in 3% low gelling temperature agarose (Sigma-Aldrich) and 80 μm sections were cut on a vibratome (VTS1000S, Leica Biosystems). Selected sections were then post-fixed in a solution of 1% OsO4 containing 1.5% potassium ferrocyanide for 60 minutes at RT and incubated overnight with 0.5% uranyl acetate 25% methanol at 4°C. Then the sections were stained with Walton’s lead aspartate for 30 minutes at 60°C, dehydrated by subsequent ethanol series with increasing concentration and finally infiltrated and embedded in resin (epon 812). Ultrathin 70nm sections were cut using an ultramicrotome (Leica Biosystems EM UC7) and collected on copper grids. Sections were imaged using a JEOL JEM1400 TEM equipped with an Olympus SIS Quemesa camera operated at 80 kV. MF-CA3 synapses were identified by their morphology, presence of multiple postsynaptic densities and high synaptic vesicle content. Overview images were taken at 2500X to analyze bouton number and area, and 5000X to analyze ultrastructure. Images were analyzed in MIB2 software (University of Helsinki) to segment the area and perimeter of individual MF-CA3 synapses, and the number and length of PSDs and AZs. Segmented images were analyzed with ImageJ software using a custom-made script. Analysis were conducted blind to genotype.

#### Lentivirus Production

For lentivirus production, HEK293T cells were transfected with mCherry or Cre-T2A-mCherry containing FUGW vector plasmids and helper plasmids PAX2 and VSVG using Fugene6 (Promega). Supernatant was collected 48 hours after transfection, spun at 2000 rpm to remove debris and filtered through a 0.22 µm filter (Millipore). 200 µl aliquots were stored at −80°C.

#### Adeno-associated virus (AAV) production and purification

HEK293T cells were seeded in DMEM (Invitrogen) containing 10% fetal bovine serum (FBS) (Invitrogen). Transfection mix, containing PEI and OptiMEM (Invitrogen), was incubated with OptiMEM containing pΔF6 helper, pAAV V2/9 Rep/Cap and pAAV-hsyn1-mGFP-T2A-Cre, pAAV-hsyn1-mGFP or pAAV-Ef1a-DIO-hChR2(E123T/T159C)-EYFP plasmids for 20 minutes at RT. Prior to adding DNA:PEI mix to HEK293T cells, DMEM-10%FBS was carefully replaced with DMEM-1%FBS. After 5 hours, DMEM-10%FBS was added to the each plate. Three days after transfection, cells were harvested, pooled, centrifuged at 1,500 rpm at 4°C for 10 minutes and cell-pellets lysed in lysis buffer (150mM NaCl and 50 mM TrisHCl-pH8.5 in endotoxin free H2O).

Lysates were frozen in dry ice and ethanol, and thawed at 37°C in a water bath, for three times. Thereafter, supernatants were collected and Benzonase (Sigma) was added to a final concentration of 50U/ml. After an incubation of 30 minutes at 37°C, lysates were centrifuged at 5000 rpm for 20 minutes at RT. Supernatant was then collected through a 0,45 μm filter (Thermo-Fisher) and carefully layered onto iodixanol gradients in 25×77mm OptiSeal tubes (Beckman Coulter). Gradients were prepared using OptiPrep iodixanol (Sigma), 5 M NaCl, 5X PBS with 1 mM MgCl2 and 2.5 mM KCl (5X PBS-MK), and sterile H2O. Gradients were centrifuged at 50,000 rpm and 12°C for 100 minutes in the Optima XE-100 Ultracentrifuge (Beckman Coulter). Next, AAVs were collected with an 18G needle (Beckman Coulter) from between the 40% and 60% layers and diluted in 5 ml 1x PBS-MK. The diluted AAVs were then desalted and concentrated by centrifugation at 5000 rpm for 30 minutes at 20°C in a pre-rinsed Amicon Ultra-15 filter (Millipore) in 1X PBS-MK. Finally, concentrated and desalted AAVs were washed by centrifugation at 5000 rpm and 20°C for 5 minutes with PBS containing 0.01% Pluronic F68 (Thermo-Fisher), aliquoted and stored at −80°C.

#### Stereotactic injections

For 3D reconstruction of MF boutons, *Igsf8* cKO mouse littermates were injected with AAV vectors expressing mGFP or mGFP-T2A-Cre in the DG at P7. For electrophysiology experiments, *Rbp4-Cre;Igsf8* cKO mouse littermates were injected with AAV vectors expressing Cre-dependent hChR2(E123T/T159C)-EYFP in the DG at P7. Prior to stereotactic injections, mice were intraperitoneally injected with 0.05 mg/kg buprenorphine (Vetergesic). After 1 hour, mice were anesthetized with 5% isoflurane and Duratears was applied to the eyes to prevent them from drying out. Mice were placed in a mouse stereotact (KOPF) equipped with a neonatal mouse adaptor (Stoelting). During the rest of the procedure 2.5% isoflurane was constantly administered. After shaving and disinfecting the mouse’s head, local anesthesia was administered by a subcutaneous injection with 100 µl lidocaine (Xylocain 1%). After 5 minutes an incision was made in the skin. Thereafter, the injector (Nanoject III Drummond) was placed at predetermined coordinates and beveled capillaries that penetrate skull and skin were used. One minute after lowering the capillary into the brain, viral mix was injected in the DG at 1 nl/s. After a 5 minute recovery, the capillary was pulled out at approximately 0.1 mm/5s. The incision was stitched with surgical glue (Millpledge Veterinary). After six hours, their health was examined and mice were injected with 0.1 mg/kg buprenorphine.

#### Image acquisition and analysis

Injected mice were perfused at P28 and brains dissected, postfixed, embedded and sectioned as before. Sections in which sparsely labelled MF axons visibly expressed GFP were immunostained with a chicken polyclonal anti-GFP primary antibody (Aves Labs) and an Alexa 488-conjugated goat-anti chicken secondary antibody (Invitrogen). Coverslips were then mounted using Mowiol 4-88 (MilliporeSigma Calbiochem). Sparsely labelled MF boutons were imaged on a Leica SP8 confocal microscope (Leica Microsystems) using a 63X oil-immersion objective and 1.75 zoom factor. Z-stacks containing the complete MF bouton structure were acquired using a z-step of 0.2 µm. Images were acquired from the CA3 hippocampal subregion. Acquired Z-stacks were then individually opened in Fiji (*55*) and registered to correct possible drift in z-axis using the StackReg plugin. In order to reconstruct individual MF boutons, aligned stacks were then uploaded in Imaris (Bitplane). The number of filopodia was manually measured in 3D and only filopodia with a minimum length of 1.5 μm were counted (*56*). The 3D reconstructions of the MF bouton core were obtained by initially segmenting the volume containing the entire MF terminal. The segmented volume was subsequently masked and the MF bouton was reconstructed using the ‘New Surface’ tool. Analysis of the volume and surface values was acquired via the ‘Statistics’ function. Morphological analyses were conducted blind to conditions.

#### Acute Slice Electrophysiology

Sagittal slices were prepared from postnatal day P27–35 *Rbp4-Cre;Igsf8* cKO littermates. Briefly, after decapitation, the brain was quickly removed and transferred into ice-cold cutting solution (in mM): 87 NaCl, 2.5 KCl, 1.25 NaH_2_PO_4_, 10 glucose, 25 NaHCO_3_, 0.5 CaCl_2_, 7 MgCl_2_, 75 sucrose, 1 kynurenic acid, 5 ascorbic acid, 3 pyruvic acid, pH 7.4 with 5% CO_2_/ 95% O_2_) and whole brain sagittal slices (300 µm) were cut using a vibratome (VT1200, Leica Biosystems). Afterwards slices were transferred to 34°C cutting solution for 45 minutes to recover and finally maintained at RT until used for recordings.

For recordings, brain slices were continuously perfused in a submerged chamber (Warner Instruments) at a rate of 3–4 ml/min with (in mM): 119 NaCl, 2.5 KCl, 1 NaH_2_PO_4_, 26 NaHCO_3_, 4 MgCl_2_, 4 M CaCl_2_, 11 glucose at pH 7.4 with 5% CO_2_/ 95% O_2_. For paired-pulse and spontaneous release recordings, 20 µM bicuculline, 100 µM AP-5 and 150 nM CNQX was added to the ACSF. Whole-cell patch-clamp recordings were done using borosilicate glass recording pipettes (resistance 3.5–5 MΩ). For paired-pulse and spontaneous release experiments we used the following internal solution (in mM): 115 CsMSF, 20 CsCl, 10 HEPES, 2.5 MgCl_2_, 4 ATP, 0.4 GTP, 10 Creatine Phosphate and 0.6 EGTA (pH 7.25). For E/I ratios we used (in mM): 132 CsMSF, 8 CsCl, 10 HEPES, 0.5 mM CaCl_2_, 1 EGTA, 10 Glucose and 5 QX-314 (pH 7.3). For experiments testing eEPSP induced action potential generation, we used the following internal medium (in mM): 135 KGluconate, 4 KCl, 2 NaCl, 10 HEPES, 4 EGTA, 4 MgATP, 0.3 NaATP (pH 7.25).

Spontaneous and evoked input to CA3 pyramidal neurons was recorded by whole-cell voltage-clamp recordings (Vm=−70 mV) from visually identifiable CA3 pyramidal neurons, using a Multiclamp 700B amplifier (Axon Instruments). Spontaneous input was analysed using Mini Analysis program (Synaptosoft), evoked data was analyzed using Clampfit 10.7 (Axon Instruments). Stereotactic injections of pAVV-Ef1a-DIO-ChR2-EYFP in the dentate gyrus of *Rbp4-Cre;Igsf8* cKO littermates at P7 resulted in expression of ChR2 only in granule cells of the dentate gyrus and therefore allowed light-induced activation specifically of the mossy fiber pathway. We used site-directed, region-controlled activation (UGA-42 Geo of Rapp OptoElectronics) of mossy fibers within stratum lucidum area close to the recorded neurons. We typically needed 2-10% laser power (0.5-2.6 mW with open aperture) for 1-2 ms, using circular area selections between 40 and 120 (32-96 µm diameter) to elicit baseline eEPSCs responses (300-400 pA). Recordings requiring significantly more optical stimulation to elicit eEPSCs were discarded. For paired-pulse ratio experiments (Vm=-70 mV), paired optical stimulations (interstimulus interval (ISI): 25, 50, 100, 200, 400, 1000 and 2000 ms) were delivered every 20 seconds (each ISI was repeated 4 times) and calculated as the eEPSC2/eEPSC1 ratio. Inhibition/excitation ratios were recorded by separating eEPSCs or eIPSCs using optical stimulation when the CA3 neuron was clamped at the reversal potential of the inhibitory (−70 mV) or excitatory (0 mV) input, respectively. Single eEPSCs and eIPSCs were repeated 8 times, every 20 seconds and averaged. For testing eEPSP induced action potential generation, we established a basal eEPSC amplitude of ∼400 pA while keeping the neuron at the chloride reversal potential (Vm=-82.6 mV). Subsequently, while recorded neurons were in current clamp (Vrest), train stimulations (10 Hz, 20 stimulations) were used to elicit mossy fiber activation to monitor CA3 pyramidal cell excitability. For all measurements we performed a minimum of three independent preparations.

### QUANTIFICATION AND STATISTICAL ANALYSIS

Data analysis was performed in GraphPad Prism 8 (GraphPad software), Clampfit 10.7 (Axon Instruments), MiniAnalysis (Synaptosoft) and Fiji (NIH). All data graphed as mean ± SEM. For all experiments, datasets were tested for normality using D’Agostino and Pearson test. If datasets passed the test, they were analyzed using Student’s unpaired t-test. Otherwise, the datasets were analyzed using nonparametric unpaired t-tests (Mann-Whitney).

## Supplemental Figures and Figure Legends

**Figure S1.**
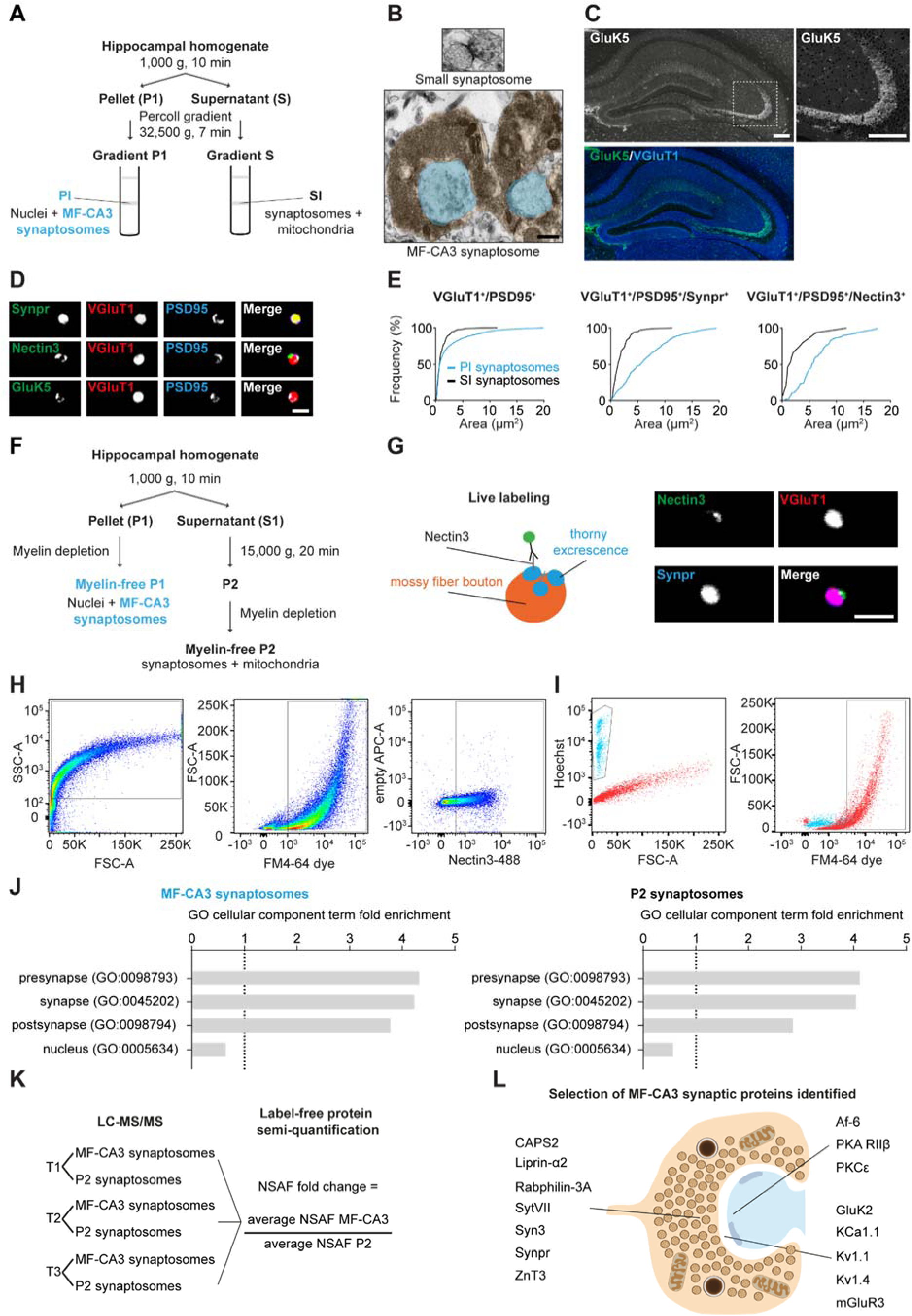
Isolation of MF-CA3 synaptosomes (Related to Figure 1) (**A**) Original method to isolate MF-CA3 synaptosomes from mouse hippocampal homogenates using Percoll gradients (*22*). (**B**) Electron microscopy image of a small synaptosome (top) and a large MF-CA3 synaptosome (bottom) detected in PI, displayed at same scale for comparison. Large MF bouton highlighted in orange filled with synaptic vesicles and containing mitochondria. Postsynaptic spines highlighted in blue. (**C**) Confocal images of P28 mouse hippocampal sections immunostained for GluK5 and VGluT1. Magnified inset of SL in CA3 is shown on the right. (**D**) Confocal images of MF-CA3 synaptosomes captured in PI immunostained for Synpr, Nectin-3, GluK5, VGluT1 and PSD95. Nectin 3 labeling was confined to small puncta as expected for a PA-localized postsynaptic protein. (**E**) Area distribution of VGluT1^+^/PSD95^+^, VGluT1^+^/PSD95^+^/Synpr^+^ and VGluT1^+^/PSD95^+^/Nectin3^+^ synaptosomes in PI and SI. (**F**) Optimized method for biochemical enrichment of MF-CA3 synaptosomes starting from P28 mouse brains. (**G**) Live labeling strategy (left) and confocal images (right) of MF-CA3 synaptosomes live-labeled with an Alexa 488-conjugated anti-Nectin-3 monoclonal antibody. (**H**) Gating strategy to sort FM4-64/Nectin3-488 double-labeled MF-CA3 synaptosomes. (**I**) Nuclei labeled with Hoechst are not labeled by FM4-64 dye. (**J**) GO analysis on all proteins detected in sorted MF-CA3 synaptosomes or P2 synaptosomes. Fold enrichment of a selection of significant cellular component terms. (**K**) Outline of the experimental set-up for the LC-MS/MS analysis of sorted MF-CA3 synaptosomes and P2 synaptosomes. NSAF FC was calculated using the average NSAF of three independent experiments. (**L**) Cartoon illustrating a selection of known MF-CA3 synaptic proteins detected in isolated MF-CA3 synaptosomes. Scale bars in (B) 0.5 μm, in (C) 200 μm, and in (D) and (G) 5 μm.

**Figure S2.**
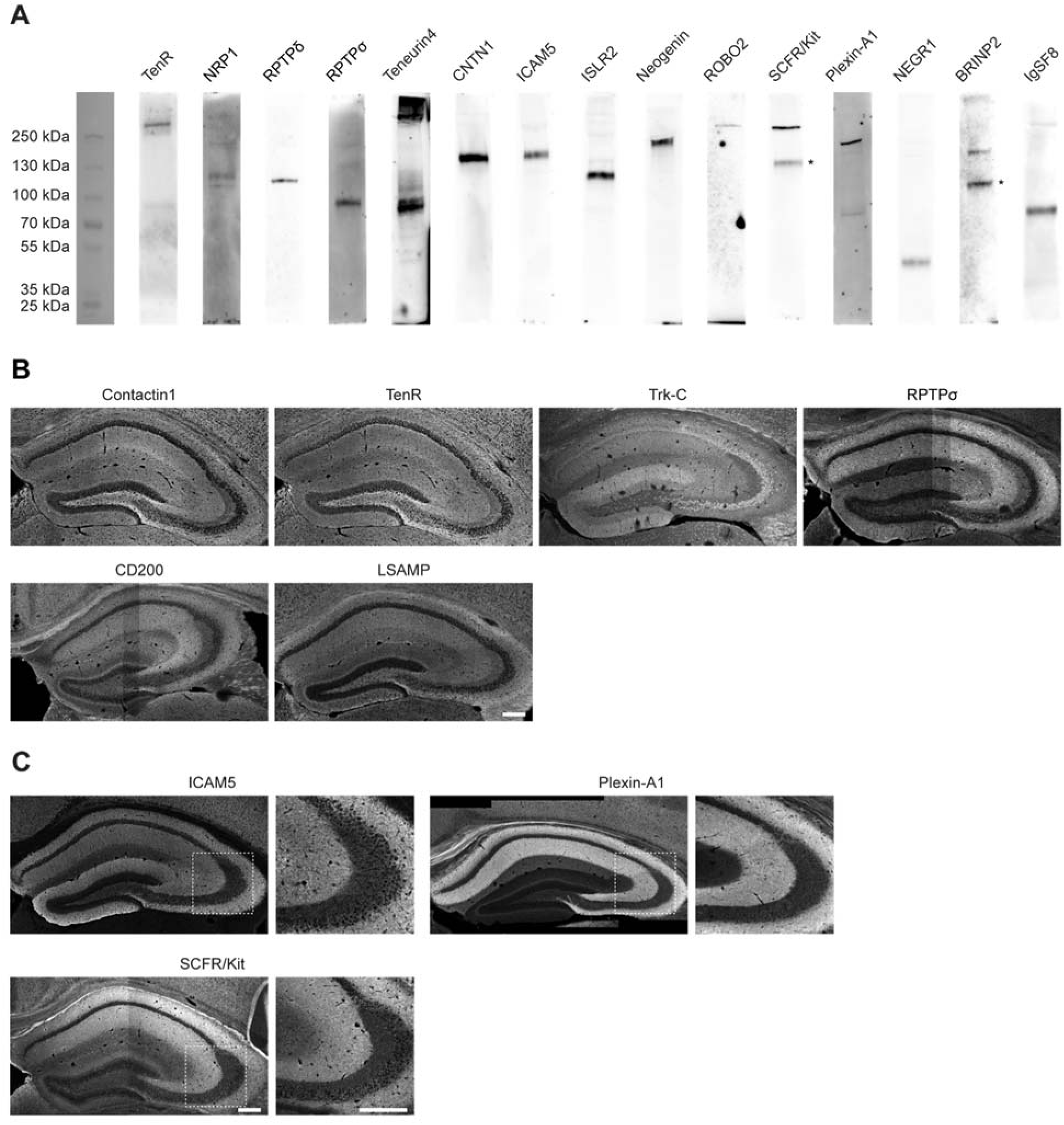
Characterization of MF-CA3 CSP composition (Related to Figure 2) (**A**) Selection of CSP antibodies working in WB using mouse hippocampal lysates. Asterisks indicate bands of expected size for respective CSPs. (**B**) Confocal images of P28 mouse hippocampal sections immunostained for 6 CSPs detected in sorted MF-CA3 synaptosomes showing broad expression in the hippocampus, including SL. (**C**) CSPs validated to be present in MF-CA3 synaptosomes by WB (Figure 2D) but with little immunoreactivity in SL. Insets show high-magnification images of the SL. Scale bars in (B) and (C) 200 μm.

**Figure S3.**
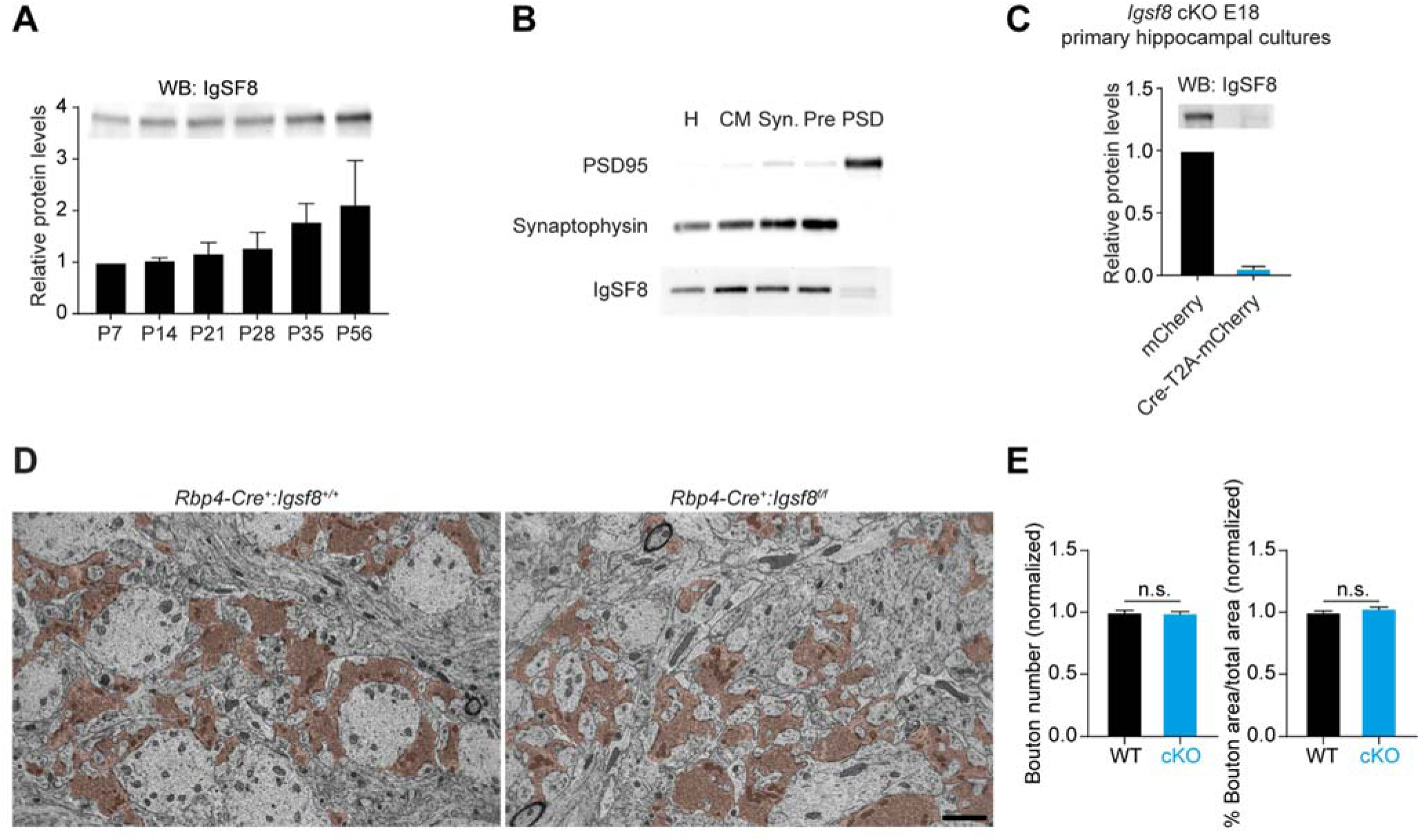
Subcellular fractionation and *Igsf8* loss of function analysis (Related to Figure 5) (**A**) Western blot analysis of IgSF8 protein expression levels in mouse hippocampal homogenates at different developmental time-points. Graph shows mean ± SEM. Quantification from two independent experiments. Total protein loading control not shown. (**B**) Subcellular fractionation using whole rat brains. H, Homogenate; CM, Crude membrane; Syn, Synaptosomes; Pre, Presynaptic fraction; PSD, Postsynaptic density fraction. (**C**) Western blot analysis of *Igsf8* cKO mouse primary hippocampal cultures infected with a lentiviral vector harboring Cre recombinase or control vector. Graph shows mean ± SEM. Quantification from three independent experiments. Total protein loading control not shown. (**D**) Electron microscope images of MF-CA3 synapses from *Rbp4-Cre*:*Igsf8* cKO and WT littermates (2500x magnification) to analyze number and area of MF boutons, highlighted in orange. (**E**) Graphs show quantification of analysis done in (D) using three littermate mice per condition (WT, n = 103 images; cKO, n = 93). Graphs show mean ± SEM. Unpaired student t-tests were used. n.s., not significant. Scale bar in (D) 2 μm.

**Figure S4.**
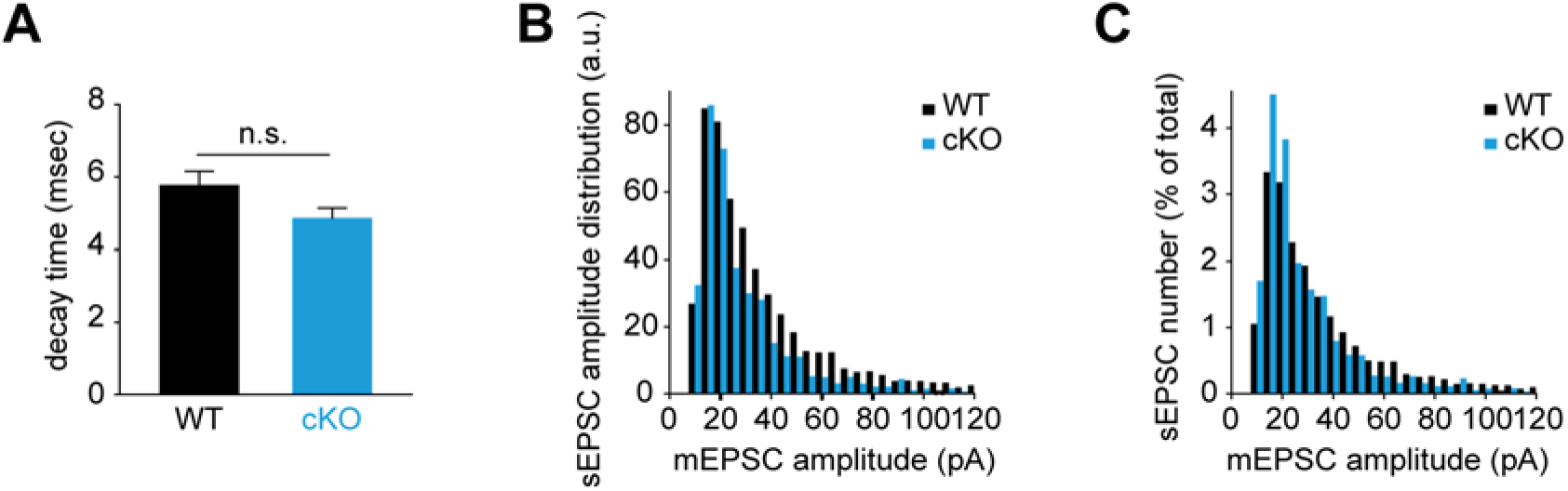
Decreased spontaneous transmission in *Igsf8* cKO CA3 neurons (Related to Figure 6) (**A**) Quantification of sEPSC decay time. (**B**) Histogram analysis of sEPSC amplitude distributions in *Rbp4-Cre:Igsf8* cKO and WT littermates. (**C**) Histogram analysis of sEPSC amplitude distributions (% of total) in *Rbp4-Cre*:*Igsf8* cKO and WT littermates. Three littermate mice were used per condition. For sEPSCs: WT, n = 31 neurons and cKO, n = 38. Graph in (A) shows mean ± SEM and Mann-Whitney test was used. n.s., not significant.

## Notes

#### Summary of Updates

New figure added that includes data on the interactome of cell-surface proteins identified in mossy fiber-CA3 synapses.

